# Inferring intestinal mucosal immune cell associated microbiome species and microbiota-derived metabolites in inflammatory bowel disease

**DOI:** 10.1101/2020.09.18.304071

**Authors:** Rajagopalan Lakshmi Narasimhan, Allison A. Throm, Jesvin Joy Koshy, Keith Metelo Raul Saldanha, Harikrishnan Chandranpillai, Rahul Deva Lal, Mausam Kumravat, Ajaya Kumar K M, Aneesh Batra, Fei Zhong, Jiajian Liu

## Abstract

Inflammatory bowel disease (IBD) is a complex, chronic inflammatory disease of the gastrointestinal tract with subtypes Crohn’s disease (CD) and ulcerative colitis (UC). While evidence indicates IBD is characterized by alterations in the composition and abundance of the intestinal microbiome, the challenge remains to specify bacterial species and their metabolites associated with IBD pathogenesis. By the integration of microbiome multi-omics data and computational methods, we provide analyses and methods for the first time to identify microbiome species and their metabolites that are associated with the human intestine mucosal immune response in patients with CD and UC at a systems level. First, we identified seven gut bacterial species and seventeen metabolites that are significantly associated with Th17 cellular differentiation and immunity in patients with active CD by comparing with those obtained in inactive CD and non-IBD controls. The seven species are *Ruminococcus gnavus, Escherichia coli, Lachnospiraceae bacterium, Clostridium hathewayi, Bacteroides faecis, Bacteroides vulgatus*, and *Akkermansia muciniphila*, and a few associated metabolites include the secondary bile acid lithocholate and three short-chain fatty acids (SCFAs): propionate, butyrate, and caproate. We next systematically characterized potential mechanistic relationships between the Th17-involved metabolites and bacterial species and further performed differential abundance analysis for both microbiome species and their metabolites in CD and UC relative to non-IBD controls with their metagenomic and metabolomic data. Based on the deconvolution of immune cell compositions from host intestinal bulk RNA-seq, we investigated changes in immune cell composition and abundance in CD and UC in comparison to non-IBD controls. Finally, we further extended our species and metabolite associations with immune cells from Th17 and Th2 cells to B cells, plasma B cells, plasmablasts, CD4+ T cells, and CD8+ T cells. While a set of associations of immune cells with bacterial species and metabolites was supported by published evidence, the new findings in this work will help to furthering our understanding of immune responses and pathogenesis in IBD.

## Introduction

Inflammatory bowel disease (IBD) is a chronic inflammatory disease of the gastrointestinal tract, which includes Crohn’s disease (CD) and ulcerative colitis (UC). CD can affect any part of gastrointestinal tract from the terminal ileum to the colon in a discontinuous pattern. In contrast, UC is localized to the colon. Over the past decade, much progress has been made including the discovery of >200 IBD associated loci^1^, identification of a collection of differentially abundant microbiome species that are associated with IBD^2–4^, and understanding of interactions of microbiota with host immune responses^5^. However, our understanding of gut microbiome dysbiosis in the pathogenesis of IBD is limited, and the etiology of IBD remains unknown.

Two important factors limit determination of the specific impact of the gut microbiota on IBD pathogenesis. First, numerous studies have demonstrated that both CD and UC are associated with reduced complexity of the commensal microbiota relative to healthy gut microbiomes, including phylum-level decreases in Firmicutes, Bacteroides, Clostridia, Lactobacillus, and Ruminococcaceae and increases in Proteobacteria and Enterobacteriaceae^6,7^. However, inconsistent results across studies and changes in a large of number gut bacteria species make it difficult to specify individual bacterial species involved in the pathogenesis of CD and UC^5,8–10^. Second, to address causal relationships between altered microbiomes and inflammation on the pathogenesis of IBD, many studies have utilized gnotobiotic mouse systems. In this system, germ-free (GF) mice are colonized with a single species or a mixture of bacteria to investigate the effects of intestinal bacteria on disease progression *in vivo*. Despite the success of GF models with individual species^8^, the challenge remains in rationally selecting species for a mixture of candidate pathogens to study the relevance of microbiota in IBD. In this work, we attempted to identify gut bacteria and their metabolites associated with human intestine mucosal immune responses at a systems level by integration of computational methods with microbiome multi-omics and host transcriptomics data^11^. Additional constraints identified for associations between intestinal immune cells with changes in gut bacteria and their metabolites will help to identify potential pathogens that are involved in intestinal inflammation and pathogenesis of IBD.

The immunological dysregulation in IBD is characterized by epithelial damage, expansion of inflammation, and infiltration of the lamina propria by many innate and adaptive immune cells that produce high levels of proinflammatory cytokines in the local tissues^12^. The role of microbiota in intestinal pathogenesis of IBD has mostly been characterized in mouse models^13^. Previous studies have focused on the roles of CD4+ helper T cell subsets and demonstrated key functions of proinflammatory Th1, Th2, and Th17 cells and anti-inflammatory regulatory T cells ^14–16^ in IBD. In recent years many studies have been extended to examine changes in T cell subsets^17–19^, plasma cells^17,20,21^, tissue macrophages, and dendritic cells^22^ to investigate pathogenesis of IBD in humans, along with the utilization of single cell RNA-seq technology to characterize changes in human intestinal cell compositions in IBD^20,23^. Evidence showed that the human intestine is massively infiltrated with plasmablasts in both CD and UC ^17,20,21^, but both CD8+ memory T cells (Trm) and gut-homing CD8+ T cells in the epithelium and lamina propria colonic tissue are reduced in CD and UC relative to controls^17^. However, the mechanism for the gut microbiome and their metabolites associating with these immune cell changes is not understood. Recently, the Huttenhower group published the most comprehensive microbiome multi-omics data from 132 patients with IBD and non-IBD controls, along with the matched host RNA-seq of human intestinal tissue biopsies^11^. This provides a valuable resource allowing us to examine how the human intestinal immune system changes with the dysbiosis of microbiota in IBD at a systems level.

Our work began with an investigation of changes in human intestinal cytokine gene expression profiles and immune cell compositions in intestinal samples by genome-wide differential expression analysis and cellular deconvolution from bulk RNA-seq data of host intestinal biopsies in patients with CD, UC, and non-IBD controls. We next identified associations of microbiome species and their metabolites with cytokine expression and immune cells. This association analysis was complemented by differential abundance analysis of microbiome species and metabolites between IBD patients and controls. By comparing the immune response-associated bacteria and their metabolites, the significant bacteria and metabolites obtained from differential abundance analysis allowed us to characterize the roles of IBD-associated changes in microbiome species and metabolites in the inflammation and pathogenesis of IBD. With this work, seven bacterial species, and several metabolites including a secondary bile acid and three short-chain fatty acids (SCFAs) were identified for their involvement in Th17 cell differentiation and immunity in active CD. Additionally, a set of bacterial species and metabolites had associations with infiltrated B cells, plasma B cells, CD4+ T cells, and CD8+ T cells in the ileum and rectum for CD and UC, respectively. Further validation of these candidate pathogen bacteria will help to improve the understanding of the pathogenesis of IBD.

## Results

### Microbiome species associated with Th17 immunity in CD patient ileum

With published RNA-seq data^11^ from ileal biopsies in CD patients and non-IBD controls and those from rectal biopsies in UC patients and non-IBD controls, we first attempted to identify the genome-wide differentially expressed genes in IBD relative to non-IBD controls. Differentially expressed genes obtained from DESeq2^24^ and edgeR^25^ were considerably consistent and revealed a total of 954 and 6177 differentially expressed genes in CD ileal and UC rectal samples, respectively (Supplemental data 1). Of these, 642 and 2790 displayed 1.5-fold-changes (Figure S1). When comparing the differentially expressed genes and 133 human cytokine genes in CytReg^26^, 42 and 44 cytokine genes were differentially expressed between CD/controls and UC/controls, respectively (Figure 1a, b; Supplemental data 2) with consistent results between edgeR and DESeq2.

**Figure 1.**
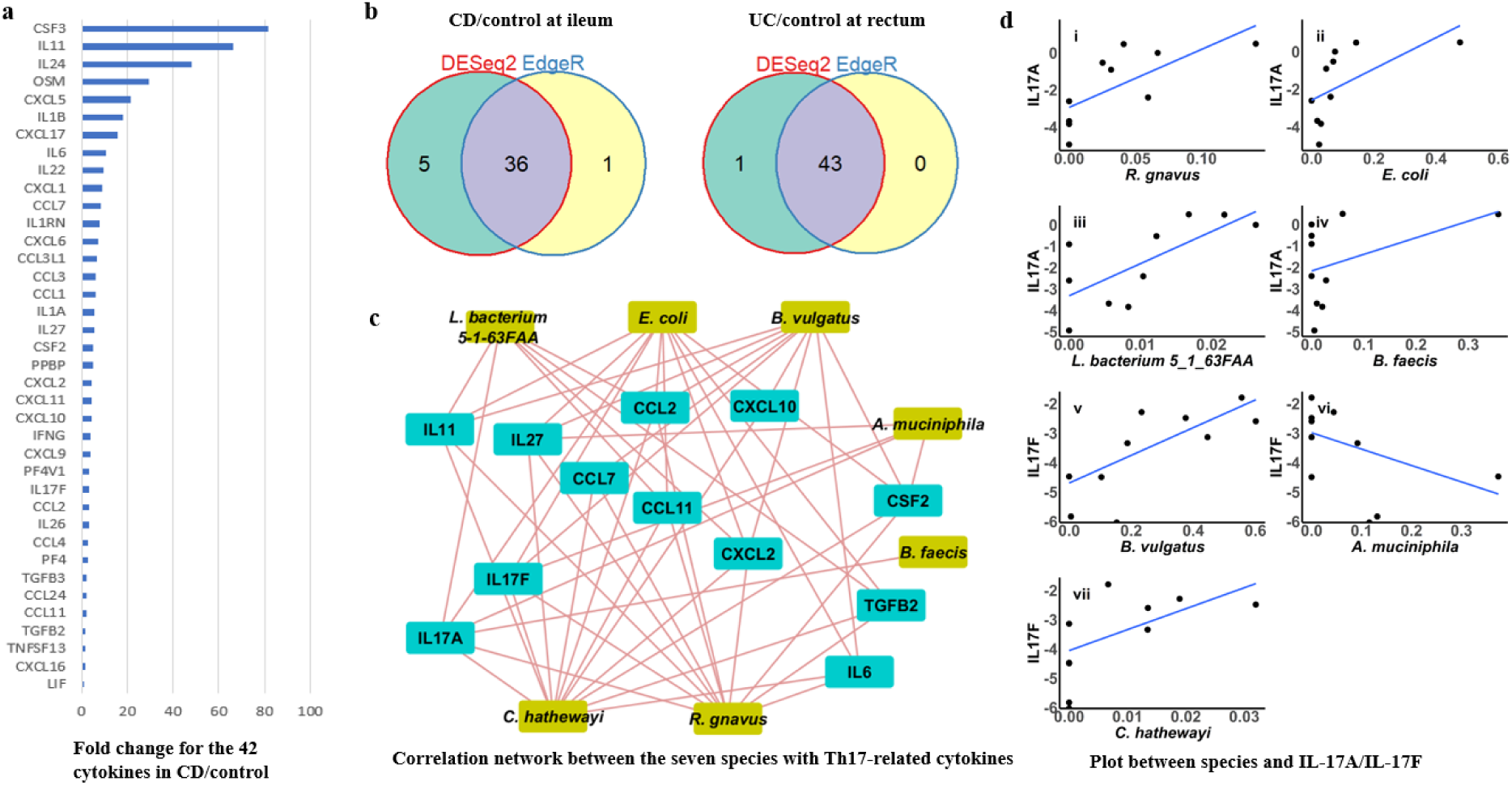
Th17 immunity associated microbiome species in active CD at ileum. **a**, Fold changes for the 42 differentially expressed cytokine genes identified in active CD relative to the non-IBD controls at ileum. **b**, Venn diagram for the numbers of differentially expressed cytokines identified by the two methods of DESeq2 and edgeR in the groups of CD/controls and UC/controls, respectively. The total number of non-redundant differential expressed cytokines identified by the two methods in CD/control and UC/control are 42 and 44, respectively. **c**, Correlation network between the seven bacterial species associated with IL-17A/Il-17F and related cytokines. Significant interaction pairs were defined based on FDR <0.05. **d**, Correlation plots between the seven bacterial species with IL-17A/ IL-17F. The seven species include *Ruminococcus gnavus, Escherichia coli, Lachnospiraceae bacterium, Clostridium hathewayi, Bacteroides faecis, Bacteroides vulgatus*, and *Akkermansia muciniphila*.

CD is a Th1 and Th17-mediated disease, whereas UC is a Th2-mediated disease^27^. Therefore, we searched for effector and signature cytokines for Th1, Th2, and Th17 immunity in the differentially expressed cytokines. Interestingly, 43% of the 42 differentially expressed cytokines identified in CD/control ileum were involved in Th17 cellular differentiation and immunity (Table S2)^28–30^. One differentially expressed cytokine, IFN-gamma, is a typical Th1-signature cytokine that is also involved in Th17 immunity. However, typical Th2 signature cytokines (IL-4, IL-5, and IL-13) were not differentially expressed between UC patients and controls. We next attempted to identify bacterial species and metabolites associated with cytokine expression, particularly in those that function in Th1 and Th17 cell immunity. We first performed pairwise correlation tests between microbial species and each of the 42 differentially expressed cytokines using two different methods, spearman rank correlation and distance covariance correlation analysis^31^. Considering that both the expression of cytokines and microbiome species in the intestine are related to disease status^17,32^, we split ileal CD samples into active and inactive groups, then performed association for patients with active CD, and further compared with those obtained from inactive CD and non-IBD controls, in which all patients were inflamed including non-IBD controls (Supplemental data 3).

In active CD, we found a total of 97 interaction pairs between 33 cytokines and 42 species (p<0.05, FDR<0.1) (Supplemental data 3). Importantly, seven of these bacterial species correlated with expression levels of Th17 effector cytokines, IL-17A and IL-17F (p <0.05, FDR <0.1). The seven associated species include *Ruminococcus gnavus, Escherichia coli, Lachnospiraceae bacterium, Clostridium hathewayi, Bacteroides faecis, Bacteroides vulgatus*, and *Akkermansia muciniphila*. All species were positively associated with IL-17A/ IL-17F except *Akkermansia muciniphila* (Figure 1c), suggesting these species are proinflammatory, while *Akkermansia muciniphila* is anti-inflammatory. IL-17A is expressed in mucosal cells and contributes to intestinal homeostasis but might promote inflammation in excess. IL-17A is mainly produced by Th17 cells. It can also be produced by γδ T cells, natural killer (NK) T cells, macrophages, neutrophils, and eosinophils^30^, but their regulation of IL-17A expression is different. For example, the production of IL-17 by γδ T cells is stimulated by IL-23 alone^33^, while IL-15 stimulates IL-17A expression in neutrophils^34^. To examine whether these bacterial species might be involved in regulation of IL-17A or IL-17F expression only or might be involved in Th17 differentiation and immunity, we performed correlation interaction network analysis between the 7 species that were found to be related to IL-17A/IL-17F expression levels and the identified associated cytokines. The interaction network includes 53 interaction edges (FDR<0.05; Figure 1d). In contrast to *Bacteroides faecis*, the other bacterial species densely interacted with multiple cytokines. Particularly, *Clostridium hathewayi, Ruminococcus gnavus*, and *Escherichia coli* significantly associated with Th17 cell differentiation inducers, including IL-6, TGFB2, IL-27, and IL-11, and cytokines dependent upon IL-17A for expression, including CSF2 and a set of IL-17 dependent chemokines involved in Th17 cellular immunity in addition to their strong associations with IL-17A and IL-17F. This suggests that these three bacterial species are most likely involved in Th17 development and immunity, while *Lachnospiraceae bacterium, Bacteroides vulgatus*, and *Akkermansia muciniphila* might also be heavily involved in Th17 development and immunity, though the they have fewer interaction edges than the three species described above. Notably, of the three most likely Th17 immunity involved species, *E. coli* strains were demonstrated to induce Th17 cell immunity in mouse models^35,36^, and *Ruminococcus gnavus* and *Clostridium hathewayi* are included in the species of 20 segmented filamentous bacteria mixture (SFB) that induced Th17 cell differentiation and immunity in a mouse model^35^.

We next examined whether our identified cytokine associated-microbiome species above are CD-specific and involved in CD progression. We first identify cytokine-associated microbiome species in inactive CD and non-IBD control ileal samples, respectively (Supplemental data 3), and then compared them with those obtained from active CD. First, in inactive CD, fewer interaction pairs between cytokines and microbiome species were detected, and no bacterial species were significantly associated with IL-17A expression, although two species, *Dorea longicatena* and *Ruminococcaceae bacterium* associated with IL-17F expression levels (p< 0.05, FDR<0.1). Second, in non-IBD controls, *Bilophila unclassified* is associated with IL-17A, while *Eubacterium ramulus* and *Lachnospiraceae bacterium* are associated with IL-17F (p<0.05, FDR<0.1). This suggests that all associated species identified in active CD are likely IBD and progression specific except for *Lachnospiraceae bacterium*. In contrast to dense interactions with Th17-invovled cytokines observed for bacterial species in active CD (Figure 1c) fewer associations were detected for species identified in inactive CD and non-IBD controls.

In this work, we cannot characterize bacterial species associated with Th1 immunity, as no single bacterial species significantly associated with interferon-gamma, a Th1-signature cytokine (p<0.05, FDR<0.1). Though 45 interaction pairs between 22 species and 23 cytokines were identified in our cytokine association analysis with species in active UC (Supplemental data 3, p<0.05, FDR<0.1), we could not derive conclusions about species-Th2 associations in active UC here, as Th2-signature cytokines, IL-4, IL-5, and IL-13 were not applied to association analysis with bacterial species because they were not in the list of 45 differentially expressed genes in UC compared to non-IBD controls, but it will be readdressed in discussion section.

### Microbiome-derived metabolites associated with Th17 immunity in CD patient ileum

The intestinal microbiota interaction with the host immune response could be directly through bacterial antigens or indirectly through their metabolites. We examined associations between cytokine expression and microbiome-derived metabolites for IBD patients using the same methods employed for the association of bacterial species with cytokine expression. In the active CD ileum, we found a total of 330 interaction pairs between 44 cytokines and 192 metabolites peaks (FDR<0.05) (Supplemental data 4). Of these, 17 metabolite features (peaks), 16 metabolites associated with IL-17A, while 1 metabolite is associated IL-17F expression (Supplemental data 4). Interestingly, 10 (66.7%) of the 17 features (peaks) associated with Th17 are bile acids and SFCAs that have been notably implicated in the pathogenesis of IBD^37,38^. Specifically, they include three SFCAs: propionate, butyrate, and caproate and five primary bile acids: cholate, chenodeoxycholate, taurocholate, alpha-muricholate, and tauro-alpha-muricholate/tauro-beta-muricholate, and 2 secondary bile acids: lithocholate and ketodeoxycholate. All three SCFAs are negatively associated with IL-17A expression levels, and a similar negative correlation pattern was observed between IL-17A expression and the secondary bile acid, lithocholate (Fig 2A-i, vii, viii, and iv). In contrast, IL-17A levels positively correlated with all five primary bile acids, along with ketodeoxycholate (Figure 2a-ii, iii, iv, v, vi). The correlation network between 18 metabolite features and 9 cytokines is shown in Figure 2b. In comparing the correlation network with analysis between bacterial species and cytokines, one important difference is that several non-Th17 pathway involved cytokines were detected, including IL-20 subfamily members IL-24 and IL-26, leukemia inhibitory factor (LIF), Th1 inducer IL-12B, and IL-11. While SCFAs primarily have edges with IL-17A, primary and secondary bile acids have more connections with other cytokine besides IL-17A and IL-17F (Figure 2b).

**Figure 2.**
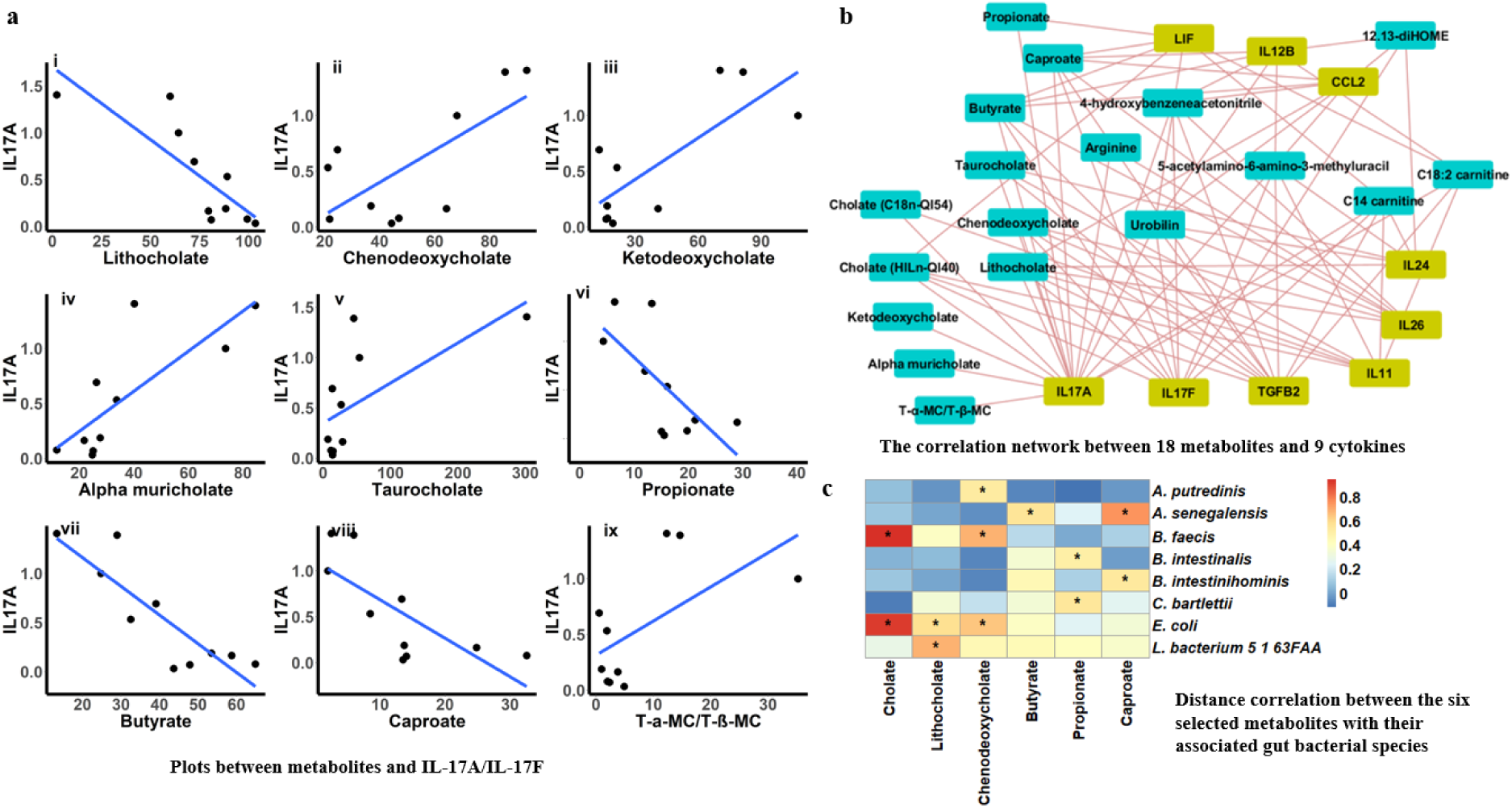
Th17 immunity associated with microbiota-derived metabolites in active CD at ileum. **a**, Correlation plots between nine metabolites and Il-17A/IL17F. These nine metabolites include lithocholate, alpha-muricholate, propionate, chenodeoxycholate,tauro-alpha-muricholate, butyrate, ketodeoxycholate, taurocholate, and caproate. **b**, Correlation network between the seven bacterial metabolites with IL-17A/Il-17F and related cytokines. Significant interaction pairs were defined based on FDR <0.05. **c**, Nonlinear distance correlations between the six selected metabolites that are associated with Th17 immunity with seven bacterial species, and the six metabolites include butyrate, caproate, propionate, cholate, chenodeoxycholate, and lithocholate. Significant interaction pairs are defined based on FDR <0.05 and are labelled with an asterisk (*).

To examine whether the IL-17A/IL-17F associated metabolites are active CD-specific, we next identified cytokine associated metabolites in inactive CD and non-IBD controls and compared with those obtained from active CD, in which all patients are inflamed including non-IBD controls. In inactive CD, there are 662 significant interactions pairs between 40 cytokines and 264 metabolites features (peaks) (FDR<0.05, Supplemental data 4), and there are 258 interaction pairs between 29 cytokines and 179 metabolites in non-IBD controls (FDR<0.05, Supplemental data 4). IL-17A correlated with 12 metabolites in inactive CD and associated with 16 metabolites in non-IBD controls. Two metabolites, taurocholate and urobilin, were found in both active CD and non-IBD controls, suggesting that tauro-conjugated primary bile acid and urobilin are not IBD-specific. However, the notable SCFAs and secondary bile acids identified in active CD are not detected in inactive CD and non-IBD. This suggest that the SFCAs propionate, butyrate, and caproate and secondary bile acid lithocholate might play unique roles in IBD disease progression and immune response.

From the 17 metabolites significantly associated with IL-17A in the active CD ileum, we chose 6 metabolites including 2 primary bile acids, 1 secondary bile acid, and 3 SCFAs to search which bacterial species in fecal microbiota are significantly associated with these metabolites in patients with active CD, inactive CD, and non-IBD controls. A total of 8, 13, and 8 bacterial species were identified in active CD, inactive CD, and non-IBD controls and associated with at least one of the six metabolites (FDR <0.05) (Supplemental data 5, Table S4), respectively. The distance correlation coefficients between the identified 8 bacterial species and the 6 Th17-immunity associated metabolites were visualized in a heatmap (Figure 2c). Of these eight species, three species, including *Escherichia coli, Lachnospiraceae bacterium*, and *Bacteroides faecis*, were also identified as Th17 immunity-associated species as described above. Specifically, *Escherichia coli* and *Lachnospiraceae bacterium* significantly associated with secondary bile acids (FDR<0.05), and *Bacteroides faecis* displayed association with primary bile acids, cholate, and chenodeoxycholate (FDR<0.05). Although the mechanism remains to be elucidated for the impact of these species on Th17 immunity, either through their antigens or their metabolites, concurrent identification of these species by two independent association approaches further supports their involvement in Th17 cellular immunity. When comparing these eight species in active CD with those in inactive CD and non-IBD controls, no pairs are conserved in inactive CD and non-IBD controls, suggesting that the metabolite associations with species are likely related to disease severity and involved in CD disease progression. However, when comparing association pairs in inactive CD with those in non-IBD controls, one widely studied butyrate-producing organism in IBD, *Faecalibacterium prausnitzii* ^39,40^ was significantly associated with butyrate (FDR <0.05) in both inactive CD and non-IBD controls, suggesting that involvement of Th17 immunity-related inflammation by *Faecalibacterium prausnitzii* might not be IBD-specific.

### IBD associated changes in the human gut microbiome and microbiome-associated metabolites at a systems level

We attempted to examine whether Th17 immunity-involved microbiome species and microbiota-derived metabolites are different in active CD relative to inactive CD and non-IBD controls. Furthermore, we attempted to identify all differentially abundant gut microbiome species and their metabolites in patients with CD and UC relative to their non-IBD controls at a systems level. Because no significantly different metagenomic species were identified previously^11^ and the sparsity of microbiome compositional data^41^, we focused on dataset stratification and methodology before performing a systematic study. First, in this analysis we only collected samples before patients were treated with drugs to minimize the impact of confounding factors on microbiome composition and abundance. Second, we extensively examined a set of quantitative methods for microbiome differential abundance analysis including the Wilcoxon rank-sum test^42^, analysis of composition of microbiomes (ANCOM)^43^, zero-inflated negative binomial distribution (ZINB)^44–47^ based methods, and zero-inflated beta-binomial (ZIBB) distribution based methods^48^. Our tests with simulated microbiomes suggest that fraction of zeros in microbiome data and means along with variance of microbiome species abundance in the non-zero fraction will affect differential abundance analysis sensitivity, specificity, and overall accuracy based on different analysis methods and data normalization methods (not published). Similar to the studies by Lloyd-Price et al.^11^, we did not identify statistically significant differentially abundant bacterial species between IBD and controls with rank-sum tests and ANCOM. However, the zero-inflated negative binomial-based method allowed us to identify differentially abundant bacterial species from patients with CD and UC patients relative to the non-IBD controls (FDR<0.05) (Supplemental data 6).

Differential abundance analysis with ZINB was performed with two datasets, one from the relative frequency data^11^ and another with geometric mean of pairwise ratios normalized (GMPR)^47^ bacterial read data generated by MetaPhlAn2^49^ (Figure 3). The two approaches are consistent, while the GMPR normalized method allowed identification of additional differentially abundant species (Figure 3a,i,ii, and iii). When comparing results in active CD with those obtained from inactive CD/non-IBD controls, five species are shared including *Bacteroides sp, Catenibacterium mitsuokai, Clostridium bolteae, Klebsiella oxytoca*, and *Klebsiella unclassified*. (Figure 3a,iv). However, 16 species were uniquely identified in active CD relative to non-IBD controls (Figure 3a). They include 8 abundance-reduced species in active CD relative to controls: *Faecalibacterium prausnitzii, Catenibacterium mitsuokai, Klebsiella pneumoniae, Clostridium ramosum, Megamonas hypermegale, Collinsella intestinalis, Parabacteroides goldsteinii, Eubacterium biforme*, and *Clostridium citroniae*, and another 8 abundance-increased species: *Bacteroides vulgatus, Sutterella wadsworthensis, Alistipes unclassified, Oscillibacter sp, Megamonas rupellensis, Streptococcus salivarius, Clostridiales bacterium*, and *Dorea formicigenerans*. ZINB calculations were also performed to detect differentially abundant species in active UC relative to non-IBD controls using the two datasets generated by the same method as described above for CD/control study (Figure 3a,iii). In total, 15 significant differentially abundant species were found from these two datasets in active UC when comparing with non-IBD controls. Of these, 4 species decreased in their abundance in active UC: *Sutterella wadsworthensis, Klebsiella pneumoniae, Veillonella parvula*, and *Clostridium ramosum*, and 11 species increased in their abundances in active UC relative to non-IBD controls: *Bacteroides sp, Citrobacter freundii, Mitsuokella unclassified, Erysipelotrichaceae bacterium, Collinsella intestinalis, Pseudomonas unclassified, Burkholderiales bacterium, Faecalibacterium prausnitzii, Collinsella unclassified, Citrobacter freundii*, and *Clostridium ramosum*.

**Figure 3.**
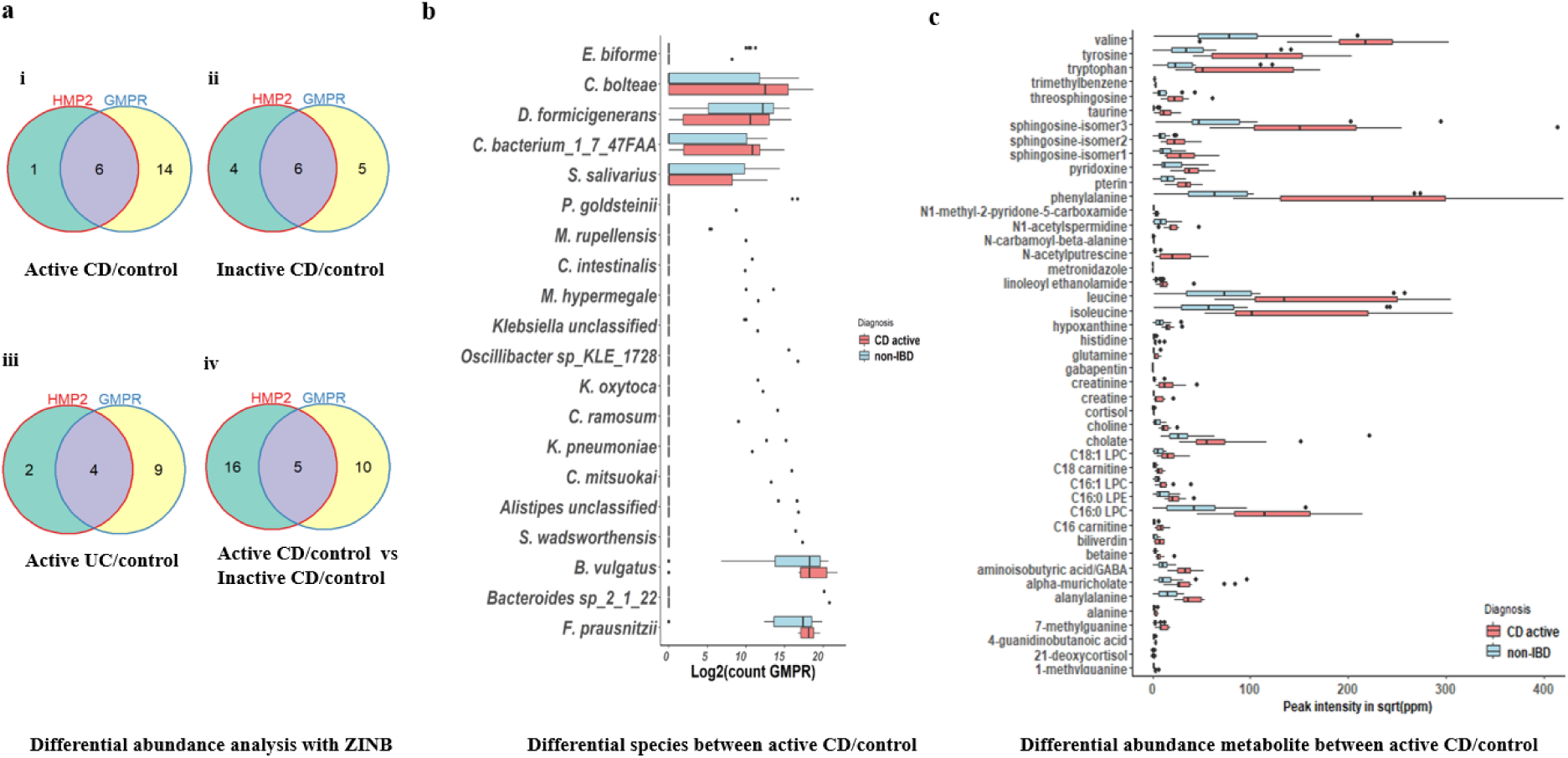
Differential abundance analysis for microbiome species and their metabolites between IBD and non-IBD controls. **a**, Venn diagram for the number of differentially abundant species between IBD and non-IBD controls using two datasets including bacterial relative frequencies and GMPR normalized bacterial read counts, respectively (from **i** to **iii). iv** is the Venn diagram for the number of differentially abundant species identified between active CD/controls and those obtained from inactive CD relative to controls. **b**, Box plots for 20 differentially abundant species in active CD relative to non-IBD controls. A large fraction of species is highly zero-inflated. **c**, Box plots for 45 differentially abundant metabolites in active CD relative to non-IBD controls.

Given the identified differentially abundant species, particularly for active CD relative to controls, we further examined which species are associated with Th17 immunity, and which are indirectly associated with Th17 immunity through their metabolites by comparison with species and metabolites associated with Th17 (Table S3). First, *Bacteroides vulgatus* is both directly associated with Th17 immunity (Figure 1c) and has a higher abundance in active CD relative to non-IBD controls. The increase of *Bacteroides vulgatus* abundance in active CD can induce more Th17 cells that in turn might worsen IBD inflammation. However, we could not rule out the critical roles of the other six species in Th17 immunity and pathogenesis in IBD, though they do not display differential abundance between active CD and non-IBD because *E. coli* was demonstrated to induce Th17 immunity^50^, while both *Bacteroides vulgatus* and *Clostridium hathewayi* are two components of 20 species that are assumed to be intestinal adherent bacteria^50^. This suggests that statistical differential abundance in active CD relative to controls might be suggestive but not necessary for Th17 immunity, particularly for analysis with limited sample sizes. Second, when we compare the list of differentially abundant species between active CD and non-IBD controls with the species that are associated with Th17-related metabolites in active CD and inactive CD, three species, *Faecalibacterium prausnitzii, Alistipes unclassified*, and *Dorea formicigenerans*, are found in common, suggesting that these species might be indirectly involved in disease progression through their Th17 immunity-related metabolites, butyrate and lithocholate.

In the metabolome data, where each sample has >80,000 measured metabolite features, we identified chemicals and chemical classes that were differentially abundant in IBD (Supplemental data 7). Forty-five differently abundant metabolites with chemical class annotation (based on HMDB) were detected in active CD relative to non-IBD controls (FDR <0.05, Figure 3c). A large fraction of these differentially abundant metabolites was previously reported by Franzosa et al^4^. Among these metabolites are α-amino acids, fatty acids, sphingolipids, and bile acids in patients with IBD. When comparing the 45 differentially abundant metabolites with the 18 metabolites significantly associated with Th17 signature cytokine expression, two metabolites, cholate and alpha-muricholate, which are primary bile acids, are in common. With limited samples, we did not find significant differences for secondary bile acids and SCFAs between active UC and non-IBD controls. Relative to a large number of differently abundant metabolites in CD in comparison with controls, only one metabolite was statistically different in UC (FDR<0.05). This may be due to approximately half of UC patients’ metabolomes being similar to non-IBD controls and small sample size^4,51^.

### Composition and variation of intestine mucosal immune cells in patients with IBD and non-IBD controls

To investigate changes of intestine mucosal immune cells in IBD patients relative to non-IBD controls, we estimated immune cell composition and abundance from bulk RNA-seq of intestinal biopsies in IBD patients using a few cellular composition deconvolution methods, including quanTIseq^52^, ABIS^53^, and xCell^54^. We first collected four independent datasets^19,55–57^, three of which are tissue samples, that provided both bulk RNA-seq data and gold-standard estimates of immune cell fractions using flow cytometry. We next performed extensive assessment on the performance of the selected methods by comparing the RNA-seq predicted results with flow cytometry measurements in the four test datasets. Each of the thee methods report cellular quantities for a collection of immune cell types. Given the limited test data, only the methods and immune cells consistently verified by different test datasets were considered for further analysis. B cell predictions by all three methods are consistent (Figure 4a). The CD4+ T cells and CD8+ T cells predicted by quanTIseq and ABIS generally meet the requirements for further analysis though the predictions for CD4+ with test dataset2 are relatively lower as are CD8+ T cell prediction with test dataset3 using quanTIseq (Figure 4a).

**Figure 4.**
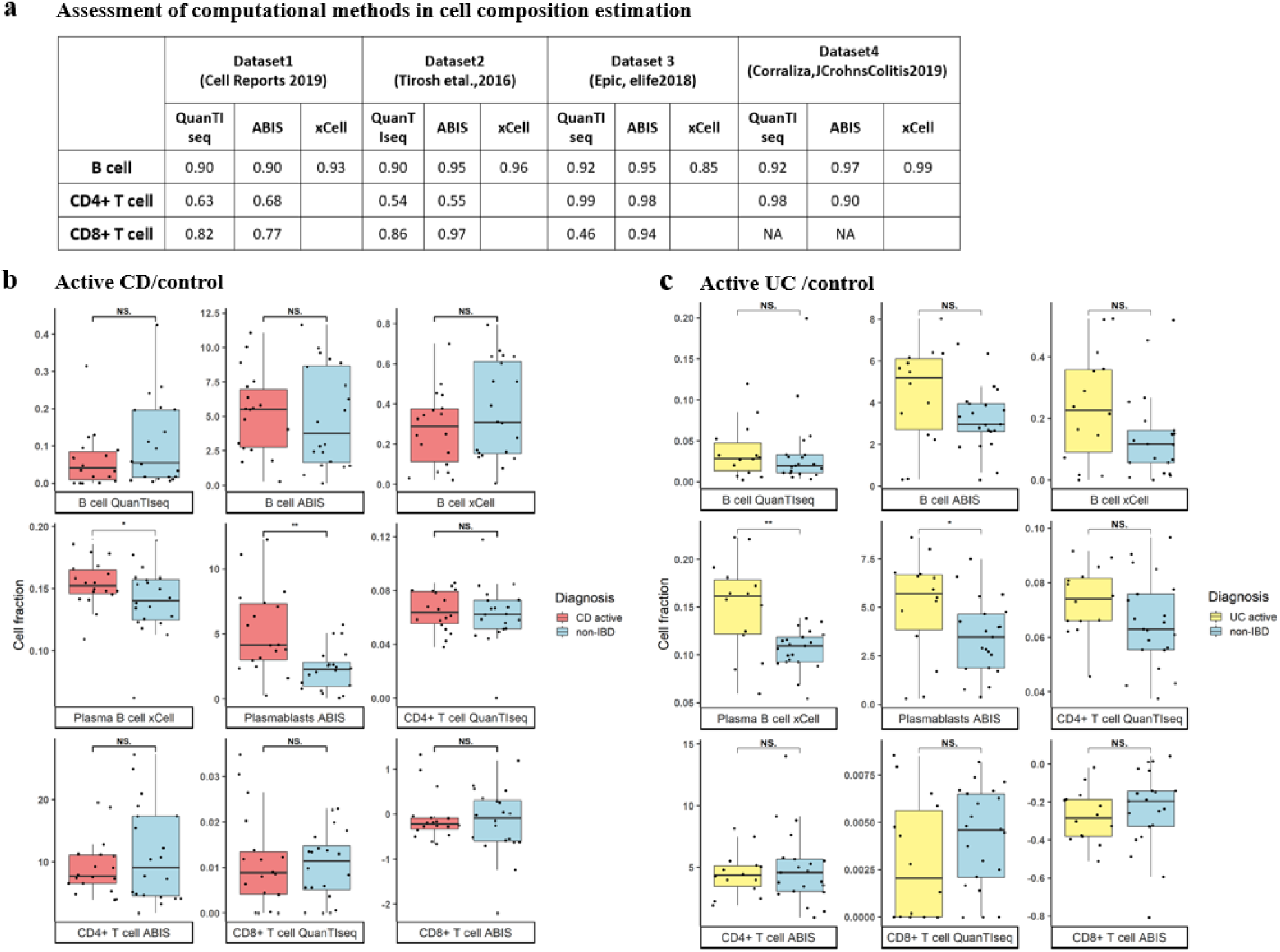
Immune cell composition estimation and comparison between IBD and non-IBD controls. **a**, Assesmsent of immune cell deconvolution methods with four indepdent datasets collected from published data^19,55–57^. Three of them are RNA-seq from tissue samples and one from peripheral blood mononuclear cells (PBMCs). **b** and **c**, Box plots for B cells, Plasma B cells, plasmablasts, CD4+ T cells, and CD8+ T cells between active CD and non-IBD controls (**b**) and between active UC and controls (**c**). Statistical tests between groups were performed with rank-sum tests. The signficantly different cells between IBD and controls are defined based on p-value <0.05 and are labelled withan asterisk (*).

B cell, plasma B cell, plasmablast, CD4+ T cell, and CD8+ T cell abundances in patients with active CD relative to non-IBD controls, and active UC relative to non-IBD controls, respectively, were visualized and statistically compared (Figure 4b, c). First, the results for B cells, plasma B cells, and plamsablast B cells in active CD/control and active UC/control comparisons indicated that they share a similar pattern (Supplemental data 8). Specifically, similar B cell levels were detected between active CD and controls and between active UC and controls. However, more infiltrated plasma B cells and plasmablasts were observed in active CD and active UC in comparison with non-IBD controls (p-value <0.05). These results are in a good agreement with previous studies with different technologies^17,20,21^. In this work, similar levels of CD8+ T cells were observed in patients with active CD or active UC relative to non-IBD controls (Supplemental data 8), and this is in concordance with a report conducted with single cell RNA-seq^20^. Similarly, no significant difference was detected in CD4+ T cell abundance between active CD/controls and active UC/controls in this work in agreement with what Noble et al. reported^17^. Interestingly we found that M1 macrophages were significantly enriched in the ileum and descending colon, while M2 macrophages show an opposing enrichment and are significantly enriched in the non-inflamed ileum. This result is consistent with the independent research by Liu et al^58^.

### Gut immune cell infiltration association with microbiome species and microbiome-associated metabolites

To determine if any other species beyond those associated with the Th17 response were associated with IBD, spearman correlation and nonlinear distance correlation were computed between immune cell frequency and microbiome species abundance for immune cell types of interest as described in previous sections, including B cells and the subsets plasmablasts and plasma B cells, CD4+ T cells, and CD8+ T cells. Microbiome species abundances positively associated with plasma B cells in the gut may be examined as potential mediators of inflammation and antibody production in IBD. In active CD patients there were 6 significant association pairs, 3 of which were for B cells or B cell subsets (FDR <0.05) (Figure 5a, S3), where none of the species were found to be differentially abundant or correlated with cytokine signatures. B cells (for both ABIS and quanTIseq deconvolutions) and *Bacteroides ovatus*, which has been used for successful monotherapy in murine colitis models^59,60^, were positively associated, suggesting that *Bacteroides ovatus* might be involved in B cell induction. Additionally, we found two negative association pairs including plasma B cells and *Eubacterium siraeum* as well as plasmablasts and *Collinsella aerofaciens*. Both *Collinsella aerofaciens* and *Bacteroides ovatus* are more coated with IgA antibodies in the gut in IBD in a previous study^61^, confirming the association of these two species with antibody production. The negative correlation could be attributed to antibacterial activities of antibodies generated by plasma B cells and plasmablast B cells or some unknown reasons. For non-IBD ileal samples, only one pair was significantly associated, B cells and *Eubacterium eligens*, which was not detected in active CD patients. While no microbiome species were associated with CD4+ T cell prevalence, CD8+ T cells were positively and negatively associated with both *Eubacterium halii* and *Anaaerostipes hadrus*, respectively, two butyrate producing bacteria; butyrate has protective effects in IBD^62–64^. In contrast to *Eubacterium halii, Anaaerostipes hadrus* was shown to increase severity of symptoms in murine colitis models^65^. CD8+ T cells were also negatively associated with *Bacteroides thetaiotaomicron*, which has been shown to promote dendritic cell responses in healthy controls but not in IBD^66^. As the role of gut CD8+ T cells in IBD is not well-studied, these three bacterial species are of interest for further studies to characterize CD8+ T cell infiltration and activity in the gut in IBD.

**Figure 5.**
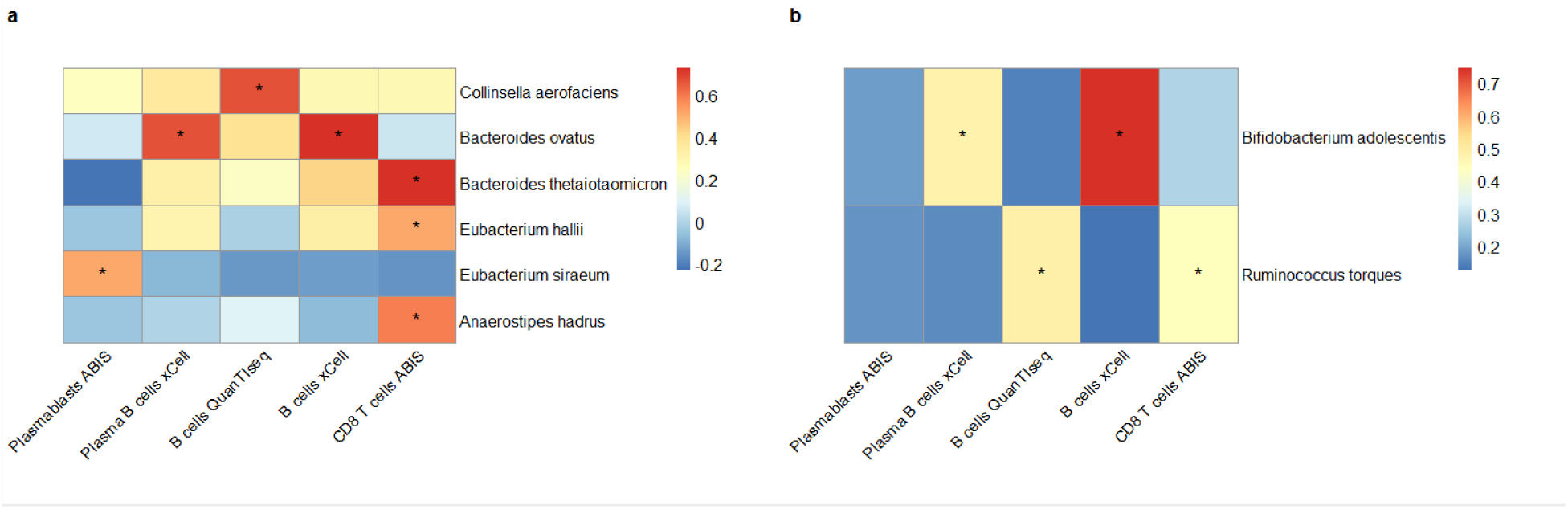
Nonlinear distance correlation coefficients between immune cells and microbes. Asterisks (*) indicate significant correlation after Benjamini-Hochberg multiple hypothesis correction for p-values < 0.05 generated by a modified t-test. **a**, CD patients with active with ileal samples, **b**, UC patients (9 active and 3 with inactive disease) with rectal samples.

For 11 UC patients, there were 4 significant pairs between immune cell frequency and microbiome species abundance, with 3 positive associations and 1 negative association (Figure 5b, S4). These species were not differentially abundant or associated with cytokine signatures. Plasmablasts were negatively associated with *Ruminococcus torques*, which has been previously demonstrated to be decreased in CD patients^67^. The decrease in plasmablasts in the gut in association with *Ruminococcus torques* levels may be due to increased differentiation of plasmablasts into antibody producing cells. B cells were positively correlated with *Bifidobacterium adolescentis* abundance (Figure 5, S4); increases in *Bifidobacterium adolescentis* in the gut in UC may lead to the recruitment of more B cells. Notably, the B cell and *Bifidobacterium adolescentis* correlation was detected for both ABIS and quanTIseq, providing further evidence for the consistency of this correlation. CD8+ T cells were positively associated with *Ruminococcus torques*. The roles of *Ruminococcus torques* and *Bifidobacterium adolescentis* in IBD are not well-studied in the literature; thus, this species may be a novel candidate to study for its effects on immune cell recruitment in IBD.

We further examined the association of all metabolites with gut immune cell infiltration in IBD to determine if other metabolites beyond those associated with Th17 response may play a role in IBD pathogenesis using a similar method as described above. For plasmablasts, a total of 19 metabolites were significant in active CD exclusively (FDR <0.05), and 9 triacylglycerol metabolites shared with ileum non-IBD controls. Whereas, no metabolites from the active CD ileum were detected in the inactive CD ileum. For plasma B cells, a total of 15 metabolites were significantly associated. Of these, one metabolite (C16:0 LPE) is significantly associated with plasma B in inactive CD ileum (Figure 6a,c), while no one association is detected in non-IBD control associated. Among these 15 metabolites are the long-chain fatty acid chemical class (myristoleate, 17-methylstearate, and heptadecanoate), one secondary bile acid (deoxycholate), one triacylglycerol (C51:0 TAG), and C16 carnitine of acyl carnitine chemical class, which were all significantly positively associated When comparing the resultant metabolites associated with plasma B cells and plasmablast B cells, it is surprising that the majority of associated metabolites for plamsablasts are involved in the glycolysis process, while most associated with plasma B cells are long-chain fatty acids. At present, we still do not know the mechanism behind it.

**Figure 6.**
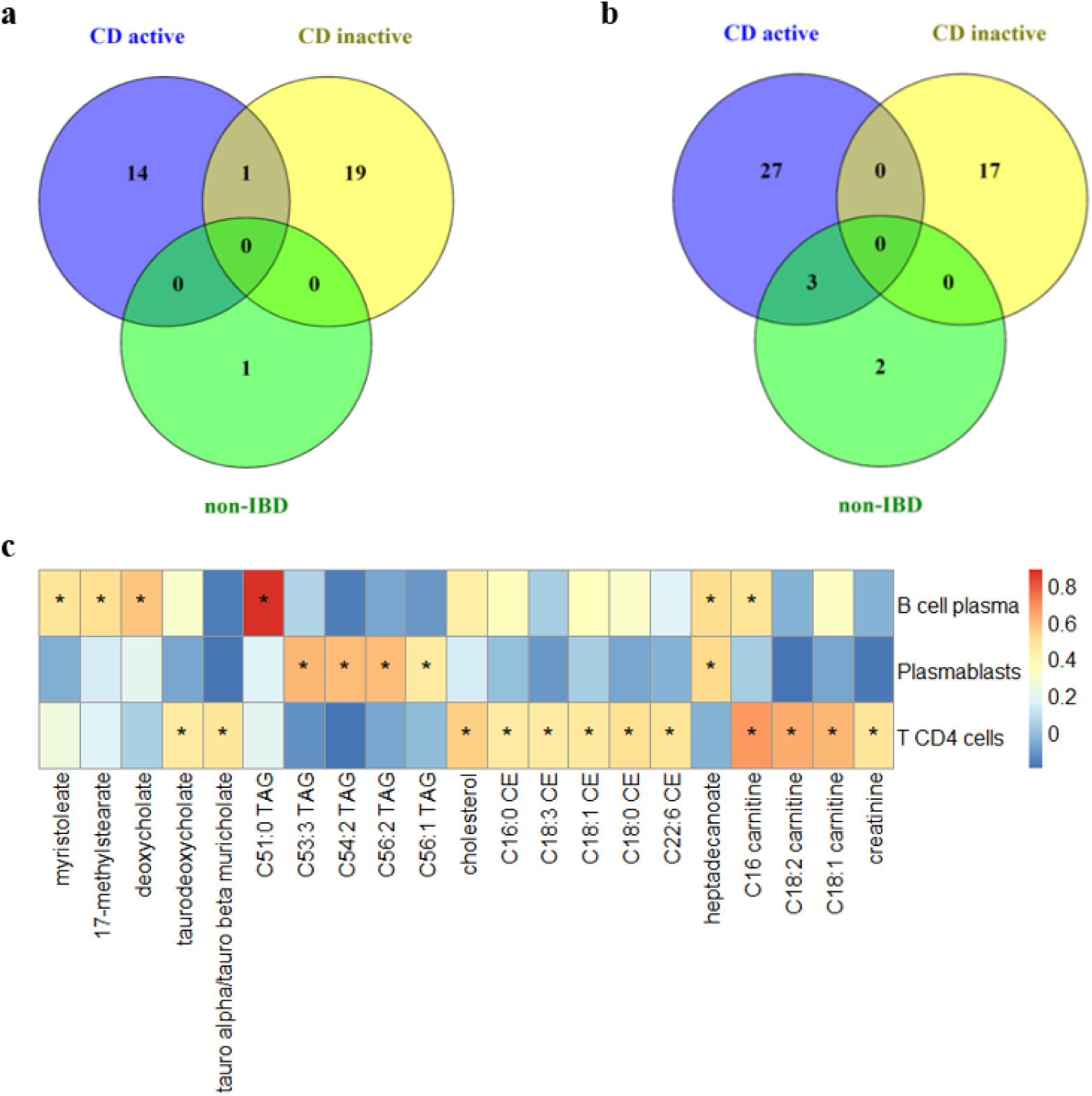
Association of immune cell infiltration with metabolites. **A**, Venn diagram of significant results for metabolite association with plasma B cells, **b**,, Venn diagram of significant results for metabolite association with CD4+ T cells. **c**, Significant metabolite association acrossileum CD active, inactive, and non-IBD controls for each cell type under study. Heatmap visualization of association between different cell types and selected metabolites based on consistent chemical group classification using distance correlation method. An asterisk (*) indicates significant interactions (FDR < 0.05). Heatmap is colored by the value of the nonlinear distance correlation coefficient.

For CD4+ T cell fractions, 30 metabolites were significant in active CD ileum, 17 in inactive CD ileum, and 5 in non-IBD ileum (Figure 6b,c). Out of these, 27 metabolites were specifically significant for active CD ileum with 24 metabolites assigned with chemical taxonomy. Five of these metabolites belong to cholesteryl esters chemical class (C16:0 CE, C18:3 CE, C18:1 CE, C18:0 CE, C22:6 CE), three acyl carnitines (C16 carnitine, C18:2 carnitine, C18:1 carnitine), two secondary bile acids (tauro alpha muricholate/tauro beta muricholate and taurodeoxycholate), and one alpha amino acid and derivative (creatinine), which have been reported to play major role in CD patients^4,38,68^. Unlike CD4+ T cells, no metabolites were found to associate with CD8+ T cells.

We further compared the identified immune cell associated significant metabolites with two sets of results: one is differential abundance metabolites, and the other is cytokine associated metabolites. First, two plasma B cells associated metabolites, C16:0 LPE and C16 carnitine, and 9 CD4+ T cell associated metabolites were all increased in active CD ileum relative toto non-IBD controls (Table S7). This suggests that an increase of these metabolites is likely involved in induction of B cell and CD4+ T cell immunity. Secondly, when we compared the significant metabolites associated with immune cells with those associated with Th17 cytokine expression, the primary bile acid tauro-alpha-muricholate/tauro-beta-muricholate and C18:2 carnitine were significantly associated with both Th17 immunity (IL-17A) and CD4+ T cells. Supplemental table 8 shows the complete list of metabolites commonly shared by Th17 immunity and immune cell types under study here. This provides additional resources to allow further characterization of connections of common metabolites with different immune responses.

## Discussion

IBD is characterized by alterations in the composition of the intestinal microbiome. Growing evidence suggests that gut dysbiosis contributes to the initiation, inflammation, and pathogenesis of IBD^6,8,10^. While these general concepts have been firmly established, specific bacterial species and their metabolites that are involved in IBD pathogenesis and their functional roles in regulating gut mucosal immune responses are not well elucidated^8^. By the integration of microbiome multi-omics data and computational methods, we provide analyses and methods for the first time, to our knowledge, to identify microbiome species and their metabolites that are statistically associated with human intestine mucosal immune responses in CD and UC at a systems level.

Our analysis was based on the dataset obtained from the Huttenhower group^11^. Their initial dataset included 132 participants, 651 biopsies, and 1785 stool samples for longitudinal IBD studies. In our analysis we only used a subset of these data, in which samples were excluded after patients were treated with antibiotic/drugs. This will help to remove the effects of medication-driven changes in the gut microbiota through tradeoff of smaller sample sizes in our work. Additionally, we further stratify our samples in our analysis based on disease status (active and inactive) and sample locations (ileum, colon, and rectum) to help to minimize the confounding factors in data analysis caused by disease types, disease severity, and tissue locations. Furthermore, all non-IBD controls in this dataset are inflamed, which provides a good control system for non-IBD inflammation in comparison to noninflamed controls. Besides the well-defined sample stratification in our study, many of our findings in this work are also attributed to the computational methods employed that improved the sensitivity of our analyses.

In this work, we first identified seven species, including *Ruminococcus gnavus, Escherichia coli, Lachnospiraceae bacterium, Clostridium hathewayi, Bacteroides faecis, Bacteroides vulgatus*, and *Akkermansia muciniphila*, and a few metabolites ranging from one secondary bile acid of lithocholate to three SCFAs of propionate, butyrate, and caproate that are significantly associated with Th17 cellular differentiation in patients with active CD when compared to inactive CD and non-IBD controls. We further detected associations of 3 species within the seven Th17 immunity-associated species including *Escherichia coli, Lachnospiraceae bacterium*, and *Bacteroides faecis*, with Th17 immunity-associated metabolites. Specifically, *Escherichia coli* and *Lachnospiraceae bacterium* were shown to significantly associate with secondary bile acids (FDR<0.05), and *Bacteroides faecis* displayed association with primary bile acids, cholate and chenodeoxycholate (FDR<0.05). This raised a few important questions as to whether an individual specific pathogen or a group of bacterial pathogens can cause IBD, whether impacts of immune response by metabolites are individual species dependent or are determined by the overall capabilities of the gut microbiota in production of these metabolites, and whether microbiome composition and abundance changes are the cause or effect of IBD. First, evidence has shown that *E. coli* strains can induce Th17 cell immunity in mouse models^35,36^, and *Clostridium hathewayi* and *Ruminococcus gnavus* are species of a segmented filamentous bacteria mixture (SFB) known to induce induced Th17 cell differentiation^35^. Our results suggest that IBD is most likely induced by more than one bacterial pathogen. Secondly, very recently Hang et al demonstrated a derivative of a secondary bile acid of lithocholic acid (LCA), 3-oxoLCA, can suppress Th17 cell differentiation by directly binding to its key transcription factor RORγt (retinoid-related orphan receptor γt) *in vitro* and *in vivo*^69^. In this work, we find that LCA and SCFs are associated with multiple species, as shown recently in LCA and butyrate^40,70^; this suggests that induction of IBD by microbiota-based metabolites is likely by the overall capabilities of the IBD-induced microbiota to produce LCA and SCFAs. Thirdly, when comparing differentially abundant species in active CD with non-IBD controls with the species that can induce Th17 immunity, a large fraction of differentially abundant microbiome species are not directly involved in induction of immune response, suggesting that the change in these bacterial abundances is likely the effect of IBD rather than the cause of IBD.

CD4+ T cells from the intestinal mucosa play a central role both in the induction and in the persistence of chronic inflammation in IBD through the production of proinflammatory cytokines^63,78^. IBD is primarily mediated by Th1 and Th17 cells in CD or Th2 cells in UC. In this work we did not find species and their metabolites associated with expression levels of the intestinal Th1 key signature cytokine, interferon-gamma, in active CD. To determine whether there are bacterial species and metabolites are associated with Th2 immunity in active UC, we further performed association again with all gut bacterial species and metabolites with key Th2 signature cytokines IL-4 and IL5, along with other Th2 cytokines IL-10, IL-13, and IL-9 that were not tested previously^14,15^. The results showed that two species, *Lachnospiraceae bacterium* and *Ruminococcus obeum* were associated with IL-4 expression levels (FDR<0.05) in active UC but not in non-IBD controls. Interestingly, *Lachnospiraceae bacterium* was also shown to be involved in Th17 immunity in active CD.

In addition to the identification of gut bacterial species and their metabolites that are associated with Th17, Th1, and Th2 immunity, we also investigated associations between microbiome species and microbiota-derived metabolites with other intestinal adaptive immune cells including B cells, B cell subsets, CD4+ T cells, and CD8+T cells. First, although two out of the three identified species that are associated with B cells or B cell subsets are detected by IgA-seq technology in IBD samples^61^, we could not absolutely claim these two species, *Bacterioides ovatus* and *Collinsella aerofaciens*, are IBD pathogenic species, as both commensal and pathogen strains can induce antigenic-specific IgA and be targeted by these antibodies once these species invade into intestine lumen after epithelial cells are broken in IBD, but they provide candidates for further characterization. Secondly, most current studies investigated the effects of metabolites on CD4+ T cells^71,72^, but less is known about their impacts on CD8+ T cell development. In this work we did not find any metabolites statistically associated with CD8+ T cells. One reason might be because these metabolites are small molecules that are not antigenic. Thirdly, while we find a few bacterial species associated with CD4+ Th17 T cells in active CD and CD4+ Th2 T cells in active UC, we did not identify any species when we attempted to associate gut bacterial species with overall CD4+ T cells. This might be attributed to multi-subset types of CD4+ T cells, each with different regulatory and action mechanisms that titrate the bacterial effect on overall CD4+ T cells. However, the species and metabolites identified in these association studies largely improve our understanding of gut immune responses in IBD.

This work provides a set of species and metabolites for further experimental verification of their roles in IBD pathogenesis. With the advance of single cell RNA-seq technology and continued refinement in quantitative methods, the association of immune responses with microbiome species and microbiota-derived metabolites can be further improved to understand the mechanism for IBD inflammation and pathogenesis.

## Methods

### Microbiome multi-omics data used in this study

We obtained the microbiome multi-omics dataset from the published results generated by the Huttenhower group in the iHMP2 project^11^. The dataset initially included 131 participants, 651 biopsies, and 1785 stool samples in their 57-week longitudinal IBD study. In order to minimize confounding effects due to medication, we only used a subset of these data that were collected prior to antibiotic/drug treatment. In this work a total of 53 IBD patients (35 CD and 18 UC) and 21 non-IBD controls were included (Table S1). All samples are inflamed including non-IBD controls. Given these patients and non-IBD controls, we collected their host RNA-seq read files generated for their ileum, colon, and rectum samples as well as their metagenome and metabolome data. For their metagenome data, we collected both their relative frequency results and raw sequencing read files separately. The summary of these data sets is described in Table S1, while sample metadata including disease severity are supplied in Supplemental data 1.

### Differential gene expression RNA-seq analysis from host intestine biopsies samples

For RNA-seq data we downloaded HMP2 raw read count data from IBDMDB^11^ and pre-processed the data by removing genes with less than 2 counts across all samples. Differential expression analysis was performed with DESeq2^24^ and edgeR^25^. Statistically significant differentially expressed genes were identified based on the Benjamini-Hochberg adjusted p-value <0.05. Over-expressed or down-regulated genes were considered with a fold change cutoff of 1.5.

To visualize differences in gene expression in CD or UC relative to their controls, the reads in each sample were normalized by counts per million reads (CPM) derived from DESeq2 and were used to conduct PCA analysis with R. To investigate gene expression patterns and signatures, we also performed hierarchical clustering analysis for gene expression, and the results were displayed with a clustering heatmap with the CRAN package pheatmap. Pathway gene set enrichment analysis was performed with the differentially expressed genes and those associated with specific biological pathways.

### Differential microbiome abundance analysis in fecal samples from CD, UC, and non-IBD controls

In this analysis two datasets were applied to identify differential microbiome species in IBD in comparison with non-IBD controls. The first data set is bacterial relative frequency results for all samples generated by the Huttenhower group^11^, and the second one was generated in this work by GMPR^47^ normalized microbiome read results. To generate the second dataset for further analysis, we first obtained metagenome sequencing read files for each sample from the IBDMDB^11^. We next employed MetaPhlAn2^49^ to identify bacterial species quantities. Instead of converting to bacterial frequencies after sample normalization, we applied the GMPR approach to normalize samples, and the quantity for each species in each sample was represented as normalized read counts.

We applied a collection of methods to identify microbiome differential abundance in this work. These methods include Wilcoxon rank-sum test^42^, analysis of composition of microbiomes (ANCOM)^43^, zero-inflated negative binomial distribution (ZINB)^44–46,73^ based methods, and zero-inflated beta-binomial (ZIBB) distribution based methods^48^. However, our differential microbiome species were primarily identified by the ZINB method. The ZINB model is a two-component mixture model combining a point mass at zero with a count distribution. Suppose that in case 1, the read count is zero, zero may come from both the point mass at zero and from the count component; in case 2, counts are generated according to the negative binomial model. the probability distribution of the ZINB random variable y_i_ can be written can be written,

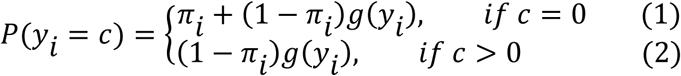

where *π*_*i*_ is the probability of subject *i* when count at zero, and *g*(*y*_*i*_) is the negative binomial function that have mean µ and variance o. The probability model was then used to calculate joint distributions across all *n* samples. In this work, our ZINB method was performed by modifying functions from the CRAN package pscl, in which we added a new method to examine model significance between the defined model and null model by the chi-square test on the loglikelihood function. ZINB models were constructed for each species. The p-value was then justified by Benjamini-Hochberg method. The species were considered statistically significant different between IBD and controls if the model justified p-value is <0.05, in which the p-value for coefficient of the non-zero-fraction is much less than 0.05.

### Differential metabolite analysis between IBD and controls in fecal samples from CD, UC, and non-IBD controls

In the metabolome data downloaded from IBDMDB, each sample contains a total of 81,867 compounds identified from four different LC-MS methods. We used a similar method as described by Franzosa et al.^4^ to perform metabolome data normalization. Specifically, the metabolome peak intensities were sum normalized within each LC-MS method. The normalized data were then used to perform Wilcoxon rank-sum tests to identify differential metabolites between IBD samples and non-IBD controls. The test p-values were justified using the Benjamini-Hochberg procedure. The metabolites were counted as statistically significant if their justified p-values were less than 0.05.

### Estimation of immune cell composition and quantities from RNA-seq of host intestinal samples

We utilized quanTIseq^52^, ABIS^55^, and xCell^54^ methods to estimate the cell-type fraction. While xCell uses a marker-genes approach with cell-type enrichment methods, quanTIseq and ABIS use a deconvolution method. Specifically, quanTIseq, ABIS, and xCell report quantities for 10, 17, and 36 types of immune cells, respectively.

We first selected four test data sets that include both RNA-seq data and immune cell quantity data determined by flow cytometry^19,55–57^. The performance of each method was assessed by comparison of their predicted results with experimental flow cytometry measurements. Given the limited experimentally measured immune cell types, our analyses were mainly focused on B cells, CD4+ T cells, and CD8+ T cells.

To estimate immune cell types in intestinal samples for CD, UC, and non-IBD controls, we used the same data sets that were used to identify genome-wide differentially expressed genes. After read QC and pretreatment by removal of bad reads and nucleotide trimming at 3’, we conducted immune cell deconvolution with the procedures as described by each of the three methods. The resultant B cells, plasma cells, plasmablasts, CD4+ T cells, and CD8+ T cells were then used for immune cell comparisons between IBD and controls, and these cells were further used for association studies with microbiome and metabolome data.

### Association of microbiome species and their metabolites with immune responses

We performed cytokine gene expression and quantified immune cells with microbiome species and their metabolites using a few methods including spearman correlation and distance correlation. Distance correlation allows for an analysis of non-linear correlation^74,75^:

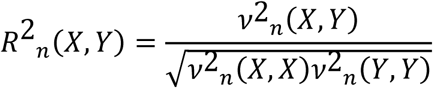

Where *R*^2^_*n*_(*X, Y*) is sample distance correlation and *v*^2^_*n*_ (*X, Y*) is distance covariance between vector X and Y.

In our association studies, the normalized read numbers from host intestinal RNA-seq were used, while species frequencies and normalized peaks were employed in cytokine association with microbiome species and their metabolites. Similarly, the same data types for microbiome and metabolome were used to compute associations between bacterial species and metabolites. In immune cell association studies, the predicted immune cell quantities were applied to associate bacterial species and metabolites. The associations between different types of data were applied for samples collected concurrently or very closely if samples were not available at the same time point. For both spearman association and distance association, a t-test was performed against null hypothesis^75^, followed by permutation testing. The initial t-test p-values were justified by the Benjamini-Hochberg procedure.

