## Supplemental appendix for "Inferring intestinal mucosal immune cell associated microbiome species and microbiota-derived metabolites in inflammatory bowel disease"

*For*

### Table of Contents

#### Supplementary Data

Supplementary data set files list……………………………………………………………..2

#### Supplementary Tables

Supplementary Table 1: Dataset summary for all data…………………..………………………..3
Supplementary Table 2: Category and functions of differentially expressed cytokines ………....4
Supplementary Table 3: IL-17 associated metabolites………………………………………5-6
Supplementary Table 4: IL-17 associated metabolites and species ………….........…………7-8
Supplementary Table 5: Differential species for CD and UC……………………..………......9-13
Supplementary Table 6: Differential metabolites for CD and UC………………………...14-15
Supplementary Table 7: Differential metabolite/immune cell comparison…………………...…16
Supplementary Table 8: Differential metabolite/cytokine comparison………………………….17

#### Supplementary Figures

Supplementary Figure 1: Differential expression analysis in CD……………………………….18
Supplementary Figure 2: Differential expression analysis in UC……………………………….19
Supplementary Figure 3: Significant microbe vs. immune cell pairs CD……………………….20
Supplementary Figure 4: Significant microbe vs. immune cell pairs UC……………………….21
Supplementary Figure 5: Metabolome and immune cell type association CD………………22-23

Supplemental References……………………………………………………………………...24

**Supplementary Data set list**

1. Genome-wide differential gene expression analysis in CD, UC, and controls with edgeR and DESeq2
2. Cytokine differential gene expression analysis in CD, UC, and controls (also for active, inactive patient subsets)
3. Association analysis between cytokine gene expression and microbiome species in CD at ileum (active /inactive), UC (active) at rectum with two methods (spearman and distance covariance correlation analysis)
4. Association analysis between cytokine gene expression with microbiome metabolites in CD (active /inactive) at ileum, UC (active) at rectum with two methods (spearman and distance covariance correlation analysis)
5. Association of metabolites with species in active CD, inactive CD and non-IBD control
6. Differential microbiome species analysis in CD at ileum, UC at rectum and control (all CD/control, active CD/control, inactive CD control, all UC, active UC/control, inactive UC/control)
7. Differential metabolites analysis in CD at ileum, UC at rectum and control (all CD/control, active CD/control, inactive CD control, all UC, active UC/control, inactive UC/control)
8. Cellular compositions in intestine samples (CD, UC and control) with methods of QuanTIseq, ABIS and xCell
9. Association analysis between predicted cell type fractions and microbiome in Ileum CD (active, inactive), non-IBD and Rectum UC active, non-IBD
10. Association analysis between predicted cell type fractions and metabolites in Ileum CD (active, inactive), non-IBD and Rectum UC active, non-IBD.
11. Metadata for RNA-seq host transcriptomics, Metagenomics and Metabolomics

**Supplemental table 1, Summary of datasets applied in this study. a**, Host RNA-seq from intestine with three locations including ileum, colon, and rectum in CD, UC, and non-IBD controls. B, Fecal metagenomic data in CD, UC, and non-IBD controls, and **c**, Fecal metabolome data in CD, UC, and non-IBD controls.

**a)**

| **Host Transcriptomics** | Ileum | Rectum | Colon |
|---|---|---|---|
| CD | 33 | 35 | 11 |
| UC | 17 | 18 | 5 |
| Non-IBD | 20 | 21 | 6 |

**b)**

| **Metagenome** | Active | Inactive | Total |
|---|---|---|---|
| CD | 10 | 8 | 18 |
| UC | 10 | 3 | 13 |
| Non-IBD | 20 | | 20 |

**c)**

| **Metabolome** | Active | Inactive | Total |
|---|---|---|---|
| CD | 11 | 8 | 19 |
| UC | 7 | 3 | 10 |
| Non-IBD | 18 | | 18 |

**Supplemental table 2, Category and functions of differentially expressed cytokines in Th17 immunity.** Forty-two cytokine genes were significantly differentially expressed between active CD and non-IBD controls. The category and gene assignment related to Th17 cellular development and immunity are assigned based on previous publications[1–5]. Out of a total 42 differentially expressed cytokines in CD relative non-IBD controls, 18 genes related to Th17 immunity, and they are labeled with red color.

| Category | Group | Cytokine genes | | functions |
|---|---|---|---|---|
| cytokine | Th17 cell development | Th17 cell inducer | IL-6, TGFB2 TGFB3 | Initiate Th17 differentiation |
| | | T17 secreted cytokine | IL-17A, IL-17F, IL-22, IFNG (interferon gamma) | Th17 cell effector cytokines |
| | IL-6 family | IL-6, IL-11, IL-27, LIF, OSM | | Pleotropic functions in inflammation |
| | IL-1 family | IL-1a, IL-1b, IL-1RN | | regulate innate and adaptive immunity |
| | IL-20 family | IL-24, IL-26 | | |
| | colony-stimulating factor | CSF2, CSF3 | | control, differentiation of monocytes, macrophages, dendritic cells (DCs), and polymorphonuclear phagocytes |
| | other | PF4V1 | | |
| Chemokine | | CXCL9 CXCL10, CXCL11, | | Neutrophil trafficking |
| | | CXCL1, CXCL2, CXCL5, CXCL6, | | T cell trafficking |
| | | CXCL16, CXCL17, CXCL9, CCL1, CCL11, CCL2, CCL24, CCL3, CCL3L1, CCL4, CCL7, | | |

**Supplemental table 3, List of IL-17A/IL17F associated microbiome-derived metabolites.** Each of the 42 differentially expressed genes in CD/controls was applied to association analysis with all metabolites with spearman rank correlation and nonlinear distance correlation analysis. This table contains the names and ID of metabolites that were found to associate with IL-17A/IL-17F (FDR <0.05). **a,** Active CD, **b,** Inactive CD, and **c,** non-IBD controls.

**a. Active CD**

| gene | compound | metabolite |
|---|---|---|
| IL17F | C18n_QI06 | 12.13-diHOME |
| IL17A | C18n_QI48 | lithocholate |
| IL17A | C18n_QI49 | chenodeoxycholate |
| IL17A | C18n_QI52 | ketodeoxycholate |
| IL17A | C18n_QI53 | alpha-muricholate |
| IL17A | C18n_QI54 | cholate |
| IL17A | C18n_QI64 | tauro-alpha-muricholate/tauro-beta-muricholate |
| IL17A | C18n_QI65 | taurocholate |
| IL17A | HILn_QI14 | 4-hydroxybenzeneacetonitrile |
| IL17A | HILn_QI28433 | Propionate |
| IL17A | HILn_QI36 | Butyrate |
| IL17A | HILn_QI37 | Caproate |
| IL17A | HILn_QI40 | Cholate |
| IL17A | HILp_QI11212 | C14 carnitine |
| IL17A | HILp_QI12459 | C18:2 carnitine |
| IL17A | HILp_QI20478 | Urobilin |
| IL17A | HILp_QI2979 | Arginine |
| IL17F | HILp_QI3985 | 5-acetylamino-6-amino-3-methyluracil |

**b. Inactive CD**

| | | |
|---|---|---|
| IL17A | HILn_QI48 | Docosapentaenoate |
| IL17A | C18n_QI42 | Docosapentaenoate |
| IL17A | C8p_QI211 | C58:10 TAG |
| IL17A | HILp_QI615 | Betaine |
| IL17A | C18n_QI36 | Eicosatrienoate |
| IL17A | C8p_QI124 | NH4_C44:2 TAG |
| IL17A | HILp_TF27 | N-acetylhistamine |
| IL17A | C8p_QI131 | NH4_C46:3 TAG |
| IL17A | C8p_QI135 | NH4_C46:1 TAG |
| IL17A | HILn_QI3 | 2-aminobutyrate |
| IL17A | HILp_QI892 | 1-methylhistamine |
| IL17A | C8p_QI85 | NH4_C22:5 CE |

**c. non-IBD control**

| | | |
|---|---|---|
| IL17A | HILn_QI65 | guanine |
| IL17A | C18n_QI40 | phytanate |
| IL17A | C8p_QI48 | linoleoylethanolamide |
| IL17A | C18n_QI65 | taurocholate |
| IL17A | C8p_QI58 | C20:0 SM |
| IL17A | HILp_QI20478 | urobilin |
| IL17A | C18n_QI61 | taurochenodeoxycholate |
| IL17A | C8p_QI212 | cholesterol |
| IL17A | C18n_QI38 | eicosenoate |
| IL17A | HILp_QI1810 | glutamate |
| IL17A | HILp_TF5 | biotin |
| IL17A | HILp_QI715 | trimethylbenzene |
| IL17A | C18n_QI33 | 17-methylstearate |
| IL17A | HILn_QI69 | homovanillate |
| IL17A | C18n_QI68 | azelate |
| IL17A | HILp_QI1048 | metformin |

**Supplemental table 4, List of microbiome species that are associated with Th17-immunity involved metabolites.** Six metabolites were selected to identify microbiome species at a systems level by association analysis with each gut bacterial species. The significant interaction pairs were defined based on FDR <0.05. The six metabolites include butyrate, caproate, propionate, cholate, chenodeoxycholate, and lithocholate. **a,** Active CD, **b,** Inactive CD, and **c,** non-IBD controls.

#### a. Active CD

| Compund | metabolite | Species/Strain |
|---|---|---|
| C18n_QI49 | chenodeoxycholate | Alistipes_putredinis\|t__GCF_000154465 |
| HILn_QI37 | caproate | Alistipes_senegalensis\|t__GCF_000312145 |
| HILn_QI36 | butyrate | Alistipes_senegalensis\|t__GCF_000312145 |
| HILn_QI40 | cholate | Bacteroides_faecis\|t__GCF_000226135 |
| C18n_QI49 | chenodeoxycholate | Bacteroides_faecis\|t__GCF_000226135 |
| HILn_QI28433 | propionate | Bacteroides_intestinalis\|t__GCF_000172175 |
| HILn_QI37 | caproate | Barnesiella_intestinihominis\|t__GCF_000296465 |
| HILn_QI28433 | propionate | Clostridium_bartlettii\|t__GCF_000154445 |
| HILn_QI40 | cholate | Escherichia_coli\|t__Escherichia_coli_unclassified |
| C18n_QI49 | chenodeoxycholate | Escherichia_coli\|t__Escherichia_coli_unclassified |
| C18n_QI48 | lithocholate | Escherichia_coli\|t__Escherichia_coli_unclassified |
| C18n_QI48 | lithocholate | Lachnospiraceae_bacterium_5_1_63FAA\|t__GCF_000185525 |

#### b. Inactive CD

| Compund | metabolite | Species/Strain |
|---|---|---|
| C18n_QI48 | lithocholate | Alistipes_unclassified |
| HILn_QI36 | butyrate | Bacteroides_caccae\|t__Bacteroides_caccae_unclassified |
| HILn_QI28433 | propionate | Bacteroides_caccae\|t__Bacteroides_caccae_unclassified |
| C18n_QI49 | chenodeoxycholate | Bacteroides_stercoris\|t__Bacteroides_stercoris_unclassified |
| C18n_QI48 | lithocholate | Bacteroides_xylanisolvens\|t__Bacteroides_xylanisolvens_unclassified |
| HILn_QI36 | butyrate | Bacteroides_xylanisolvens\|t__Bacteroides_xylanisolvens_unclassified |
| HILn_QI28433 | propionate | Bacteroides_xylanisolvens\|t__Bacteroides_xylanisolvens_unclassified |
| HILn_QI37 | caproate | Bilophila_unclassified |
| C18n_QI48 | lithocholate | Dialister_invisus\|t__GCF_000160055 |
| HILn_QI36 | butyrate | Dialister_invisus\|t__GCF_000160055 |
| HILn_QI28433 | propionate | Dialister_invisus\|t__GCF_000160055 |

| | | |
|---|---|---|
| C18n_QI48 | lithocholate | Dorea_formicigenerans\|t__Dorea_formicigenerans_unclassified |
| HILn_QI36 | butyrate | Faecalibacterium_prausnitzii\|t__Faecalibacterium_prausnitzii_unclassified |
| HILn_QI28433 | propionate | Haemophilus_parainfluenzae\|t__Haemophilus_parainfluenzae_unclassified |
| C18n_QI48 | lithocholate | Lachnospiraceae_bacterium_3_1_46FAA\|t__GCF_000209405 |
| HILn_QI28433 | propionate | Lachnospiraceae_bacterium_3_1_46FAA\|t__GCF_000209405 |
| HILn_QI28433 | propionate | Prevotella_copri\|t__GCF_000157935 |
| C18n_QI49 | chenodeoxycholate | Ruminococcus_gnavus\|t__Ruminococcus_gnavus_unclassified |
| HILn_QI37 | caproate | Ruminococcus_torques\|t__Ruminococcus_torques_unclassified |

#### c. non-IBD control

| **Compound** | **metabolite** | **Species/Strain** |
|---|---|---|
| HILn_QI37 | caproate | Bacteroides_finegoldii\|t__Bacteroides_finegoldii_unclassified |
| HILn_QI36 | butyrate | Faecalibacterium_prausnitzii\|t__Faecalibacterium_prausnitzii_unclassified |
| HILn_QI28433 | propionate | Lachnospiraceae_bacterium_3_1_57FAA_CT1\|t__GCF_000218405 |
| HILn_QI37 | caproate | Prevotella_copri\|t__GCF_000157935 |
| HILn_QI37 | caproate | Roseburia_hominis\|t__GCF_000225345 |
| C18n_QI48 | lithocholate | Streptococcus_salivarius\|t__Streptococcus_salivarius_unclassified |
| C18n_QI48 | lithocholate | Veillonella_dispar\|t__GCF_000160015 |
| C18n_QI48 | lithocholate | Veillonella_parvula\|t__Veillonella_parvula_unclassified |

**Supplemental table 5, Differential species for CD/controls (all, active, inactive), UC/controls (active)**

List of significant species and from the differential analysis of disease vs control using zero inflated negative binomial (ZINB) from HMP2 and GMPR normalized data. The overlap column represents the species overlap between the two sets. **a,** All CD/non-IBD, **b,** Active CD/non-IBD, **c,** Inactive CD/non-IBD **d,** All UC/non-IBD and, **e,** Active UC/non-IBD.

**a) All CD/non-IBD**

| HMP2 | GMPR normalized | Overlap |
|---|---|---|
| Porphyromonas_asaccharolytica\|t__Porphyromonas_asaccharolytica_unclassified | Citrobacter_freundii\|t__Citrobacter_freundii_unclassified | Porphyromonas_asaccharolytica\|t__Porphyromonas_asaccharolytica_unclassified |
| Clostridium_ramosum\|t__GCF_000154485 | Sutterella_wadsworthensis\|t__GCF_000297775 | Clostridium_ramosum\|t__GCF_000154485 |
| Oscillibacter_sp_KLE_1728\|t__GCF_000469425 | Bacteroides_vulgatus\|t__Bacteroides_vulgatus_unclassified | Oscillibacter_sp_KLE_1728\|t__GCF_000469425 |
| Porphyromonas_uenonis\|t__GCF_000174775 | Clostridium_ramosum\|t__GCF_000154485 | Porphyromonas_uenonis\|t__GCF_000174775 |
| Ruminococcus_torques\|t__Ruminococcus_torques_unclassified | Porphyromonas_asaccharolytica\|t__Porphyromonas_asaccharolytica_unclassified | Ruminococcus_torques\|t__Ruminococcus_torques_unclassified |
| Citrobacter_freundii\|t__Citrobacter_freundii_unclassified | Ruminococcus_torques\|t__Ruminococcus_torques_unclassified | Citrobacter_freundii\|t__Citrobacter_freundii_unclassified |
| Clostridium_bartlettii\|t__GCF_000154445 | Subdoligranulum_unclassified | Clostridium_bartlettii\|t__GCF_000154445 |
| | Oscillibacter_sp_KLE_1728\|t__GCF_000469425 | |
| | Klebsiella_unclassified | |
| | Megamonas_hypermegale\|t__GCF_000209975 | |
| | Collinsella_intestinalis\|t__GCF_000156175 | |
| | Megamonas_rupellensis\|t__GCF_000378365 | |
| | Bacteroides_uniformis\|t__Bacteroides_uniformis_unclassified | |
| | Dorea_longicatena\|t__GCF_000154065 | |
| | Bacteroides_caccae\|t__Bacteroides_caccae_unclassified | |
| | Veillonella_parvula\|t__Veillonella_parvula_unclassified | |
| | Clostridium_hathewayi\|t__Clostridium_hathewayi_unclassified | |
| | Eubacterium_biforme\|t__GCF_000156655 | |

| | |
|---|---|
| | Bifidobacterium_longum\|t__Bifidobacterium_longum_unclassified |
| | Porphyromonas_uenonis\|t__GCF_000174775 |
| | Clostridium_leptum\|t__GCF_000154345 |
| | Alistipes_shahii\|t__GCF_000210575 |
| | Bacteroides_clarus\|t__GCF_000195615 |
| | Clostridium_bartlettii\|t__GCF_000154445 |

### b) Active CD/non-IBD

| HMP2 | GMPR normalized | Overlap |
|---|---|---|
| Faecalibacterium_prausnitzii\|t__Faecalibacterium_prausnitzii_unclassified | Faecalibacterium_prausnitzii\|t__Faecalibacterium_prausnitzii_unclassified | Faecalibacterium_prausnitzii\|t__Faecalibacterium_prausnitzii_unclassified |
| Bacteroides_sp_2_1_22\|t__GCF_000162155 | Bacteroides_sp_2_1_22\|t__GCF_000162155 | Bacteroides_sp_2_1_22\|t__GCF_000162155 |
| Catenibacterium_mitsuokai\|t__GCF_000173795 | Bacteroides_vulgatus\|t__Bacteroides_vulgatus_unclassified | Catenibacterium_mitsuokai\|t__GCF_000173795 |
| Clostridium_ramosum\|t__GCF_000154485 | Sutterella_wadsworthensis\|t__GCF_000297775 | Clostridium_ramosum\|t__GCF_000154485 |
| Oscillibacter_sp_KLE_1728\|t__GCF_000469425 | Alistipes_unclassified | Oscillibacter_sp_KLE_1728\|t__GCF_000469425 |
| Klebsiella_unclassified | Catenibacterium_mitsuokai\|t__GCF_000173795 | Klebsiella_unclassified |
| Clostridium_citroniae\|t__GCF_000233455 | Klebsiella_pneumoniae\|t__Klebsiella_pneumoniae_unclassified | |
| | Clostridium_ramosum\|t__GCF_000154485 | |
| | Klebsiella_oxytoca\|t__Klebsiella_oxytoca_unclassified | |
| | Oscillibacter_sp_KLE_1728\|t__GCF_000469425 | |
| | Klebsiella_unclassified | |
| | Megamonas_hypermegale\|t__GCF_000209975 | |

| | |
|---|---|
| | Collinsella_intestinalis\|t__GCF_000156175 |
| | Megamonas_rupellensis\|t__GCF_000378365 |
| | Parabacteroides_goldsteinii\|t__Parabacteroides_goldsteinii_unclassified |
| | Streptococcus_salivarius\|t__Streptococcus_salivarius_unclassified |
| | Clostridiales_bacterium_1_7_47FAA\|t__GCF_000155435 |
| | Dorea_formicigenerans\|t__Dorea_formicigenerans_unclassified |
| | Clostridium_bolteae\|t__Clostridium_bolteae_unclassified |
| | Eubacterium_biforme\|t__GCF_000156655 |

### c) Inactive CD/non-IBD

| HMP2 | GMPR normalized | Overlap |
|---|---|---|
| Bacteroides_sp_2_1_22\|t__GCF_000162155 | Bacteroides_sp_2_1_22\|t__GCF_000162155 | Bacteroides_sp_2_1_22\|t__GCF_000162155 |
| Porphyromonas_asaccharolytica\|t__Porphyromonas_asaccharolytica_unclassified | Citrobacter_freundii\|t__Citrobacter_freundii_unclassified | Porphyromonas_asaccharolytica\|t__Porphyromonas_asaccharolytica_unclassified |
| Catenibacterium_mitsuokai\|t__GCF_000173795 | Catenibacterium_mitsuokai\|t__GCF_000173795 | Catenibacterium_mitsuokai\|t__GCF_000173795 |
| Desulfovibrio_piger\|t__GCF_000156375 | Parabacteroides_johnsonii\|t__Parabacteroides_johnsonii_unclassified | Desulfovibrio_piger\|t__GCF_000156375 |
| Porphyromonas_uenonis\|t__GCF_000174775 | Porphyromonas_asaccharolytica\|t__Porphyromonas_asaccharolytica_unclassified | Porphyromonas_uenonis\|t__GCF_000174775 |
| Bifidobacterium_pseudocatenulatum\|t__GCF_000173435 | Desulfovibrio_piger\|t__GCF_000156375 | Citrobacter_freundii\|t__Citrobacter_freundii_unclassified |
| Citrobacter_freundii\|t__Citrobacter_freundii_unclassified | Clostridium_bolteae\|t__GCF_000154365 | |

| | |
|---|---|
| Eubacterium_siraeum\|t__Eubacterium_siraeum_unclassified | Bacteroides_caccae\|t__Bacteroides_caccae_unclassified |
| Klebsiella_oxytoca\|t__Klebsiella_oxytoca_unclassified | Bacteroides_clarus\|t__GCF_000195615 |
| Klebsiella_unclassified | Ruminococcus_torques\|t__Ruminococcus_torques_unclassified |
| | Porphyromonas_uenonis\|t__GCF_000174775 |

#### d) All UC/non-IBD

| HMP2 | GMPR normalized | Overlap |
|---|---|---|
| Bacteroides_sp_4_3_47FAA\|t__GCF_000158515 | Bacteroides_sp_4_3_47FAA\|t__GCF_000158515 | Bacteroides_sp_4_3_47FAA\|t__GCF_000158515 |
| Clostridium_ramosum\|t__GCF_000154485 | Citrobacter_freundii\|t__Citrobacter_freundii_unclassified | Mitsuokella_unclassified |
| Mitsuokella_unclassified | Sutterella_wadsworthensis\|t__GCF_000297775 | Citrobacter_freundii\|t__Citrobacter_freundii_unclassified |
| Citrobacter_freundii\|t__Citrobacter_freundii_unclassified | Klebsiella_pneumoniae\|t__Klebsiella_pneumoniae_unclassified | |
| Oscillibacter_sp_KLE_1728\|t__GCF_000469425 | Mitsuokella_unclassified | |
| Peptostreptococcaceae_noname_unclassified | Erysipelotrichaceae_bacterium_6_1_45\|t__GCF_000242175 | |
| | Collinsella_intestinalis\|t__GCF_000156175 | |
| | Pseudomonas_unclassified | |
| | Ruminococcus_torques\|t__Ruminococcus_torques_unclassified | |

#### e) Active UC/non-IBD

| HMP2 | GMPR normalized | Overlap |
|---|---|---|
| Porphyromonas_asaccharolytica\|t__Porphyromonas_asaccharolytica_unclassified | Bacteroides_sp_4_3_47FAA\|t__GCF_000158515 | Mitsuokella_unclassified |
| Porphyromonas_uenonis\|t__GCF_000174775 | Citrobacter_freundii\|t__Citrobacter_freundii_unclassified | Bacteroides_sp_4_3_47FAA\|t__GCF_000158515 |

| | | |
|---|---|---|
| Mitsuokella_unclassified | Sutterella_wadsworthensis\|t__GCF_000297775 | Citrobacter_freundii\|t__Citrobacter_freundii_unclassified |
| Bacteroides_sp_4_3_47FAA\|t__GCF_000158515 | Klebsiella_pneumoniae\|t__Klebsiella_pneumoniae_unclassified | Clostridium_ramosum\|t__GCF_000154485 |
| Citrobacter_freundii\|t__Citrobacter_freundii_unclassified | Mitsuokella_unclassified | |
| Clostridium_ramosum\|t__GCF_000154485 | Erysipelotrichaceae_bacterium_6_1_45\|t__GCF_000242175 | |
| | Collinsella_intestinalis\|t__GCF_000156175 | |
| | Pseudomonas_unclassified | |
| | Veillonella_parvula\|t__Veillonella_parvula_unclassified | |
| | Burkholderiales_bacterium_1_1_47\|t__GCF_000144975 | |
| | Faecalibacterium_prausnitzii\|t__Faecalibacterium_prausnitzii_unclassified | |
| | Clostridium_ramosum\|t__GCF_000154485 | |
| | Collinsella_unclassified | |

**Supplemental table 6, Differential metabolites for active CD/non-IBD and UC/non-IBD (all, active).**
List of significant metabolites along with compound ID, from the differential analysis of disease vs control using the Wilcoxon rank sum test. The significant metabolites were defined based on Benjamini-Hochberg FDR corrected p-value < 0.05. **a,** Active CD/non-IBD, **b,** All UC/non-IBD, and **c,** Active UC/non-IBD

a. **Active CD/non-IBD**

| **Compound** | **Metabolite** | **FDR p-value** |
|---|---|---|
| HILp_QI522 | creatinine | 0.00248 |
| HILp_QI1093 | N-acetylputrescine | 0.00248 |
| HILp_QI11887 | C16 carnitine | 0.00275 |
| HILp_TF28 | aminoisobutyric acid/GABA | 0.00279 |
| HILp_QI1110 | creatine | 0.00308 |
| HILp_TF36 | alanylalanine | 0.00396 |
| HILp_QI2014 | N1-methyl-2-pyridone-5-carboxamide | 0.00451 |
| HILp_QI12539 | C18 carnitine | 0.00607 |
| HILp_TF58 | valine | 0.00653 |
| HILp_QI868 | taurine | 0.00653 |
| HILp_QI1144 | N-carbamoyl-beta-alanine | 0.00653 |
| HILp_QI2162 | histidine | 0.0082 |
| HILp_QI615 | betaine | 0.0089 |
| HILp_QI10472 | 21-deoxycortisol | 0.0089 |
| HILp_QI2601 | 7-methylguanine | 0.00992 |
| HILp_TF55 | phenylalanine | 0.01254 |
| HILp_QI2602 | 1-methylguanine | 0.01254 |
| HILp_QI9633 | linoleoyl ethanolamide | 0.01254 |
| HILp_QI331 | choline | 0.01587 |
| HILp_QI3549 | N1-acetylspermidine | 0.01587 |
| HILp_TF15 | pyridoxine | 0.01587 |
| HILp_TF57 | tyrosine | 0.01759 |
| HILp_TF30 | C16:0 LPC | 0.01759 |
| C8p_QI03 | C16:1 LPC | 0.01927 |
| HILp_QI10949 | cortisol | 0.01927 |
| HILp_TF7 | pterin | 0.01927 |
| HILp_QI8646 | threosphingosine | 0.02111 |
| HILp_TF12 | sphingosine-isomer3 | 0.02111 |
| HILp_TF56 | tryptophan | 0.02318 |
| HILp_QI168 | alanine | 0.02531 |
| HILp_QI715 | trimethylbenzene | 0.02763 |
| HILp_QI15155 | C18:1 LPC | 0.02763 |
| HILp_TF3 | sphingosine-isomer2 | 0.02763 |

| | | |
|---|---|---|
| HILp_QI1303 | hypoxanthine | 0.03023 |
| HILp_QI1788 | glutamine | 0.03023 |
| HILp_TF2 | sphingosine-isomer1 | 0.03023 |
| C18n_QI53 | alpha-muricholate | 0.03305 |
| HILp_QI1750 | 4-guanidinobutanoic acid | 0.03305 |
| HILp_QI19408 | biliverdin | 0.03305 |
| HILp_QI2874 | gabapentin | 0.03305 |
| HILp_QI2850 | metronidazole | 0.03305 |
| HILp_TF54 | leucine | 0.03605 |
| HILp_TF53 | isoleucine | 0.03605 |
| HILp_QI13155 | C16:0 LPE | 0.03605 |
| HILn_QI40 | cholate | 0.04302 |

**b. All UC/non-IBD**

| **Compound** | **Metabolite** | **FDR p-value** |
|---|---|---|
| HILp_QI2014 | N1-methyl-2-pyridone-5-carboxamide | 0.01144 |
| HILn_QI93 | orotate | 0.02836 |

**c. Active UC/non-IBD**

| **Compound** | **Metabolite** | **FDR p-value** |
|---|---|---|
| HILn_QI93 | orotate | 0.04347 |

**Supplemental table 7, Comparison of significant metabolites from association analysis and differential analysis for ileum CD active**

List of all significant metabolites shared between the metabolite association analysis with different cell types under study and metabolite differential analysis.

| Compound | Metabolite name | Cell type |
|---|---|---|
| HILp_QI13155 | C16:0 LPE | B Plasma cells |
| HILp_QI11887 | C16 carnitine | B Plasma cells |
| HILp_QI10472 | 21-deoxycortisol | CD4+ T cells |
| HILp_QI11887 | C16 carnitine | CD4+ T cells |
| HILp_QI10949 | cortisol | CD4+ T cells |
| HILp_QI522 | creatinine | CD4+ T cells |
| HILp_QI9633 | linoleoyl ethanolamide | CD4+ T cells |
| HILp_QI2850 | metronidazole | CD4+ T cells |
| HILp_TF2 | sphingosine-isomer1 | CD4+ T cells |
| HILp_TF3 | sphingosine-isomer2 | CD4+ T cells |
| HILp_QI868 | taurine | CD4+ T cells |

**Supplemental table 8, Comparison of significant metabolites from association analysis with cell types and cytokine/chemokine gene for ileum CD active**

List of all significant metabolites shared between the metabolite association analysis with different cytokine/chemokine genes and association analysis with different cell types under study.

| Gene | Compound | Metabolite name | Immune cell type |
|---|---|---|---|
| CXCL17 | C8p_QI82 | C22:6 CE | CD4+ T cells |
| CXCL17 | HILn_TF16 | suberate | CD4+ T cells |
| IL13 | HILn_QI11 | 3-methylglutaconate | Plasmablasts |
| IL17A | HILp_QI12459 | C18:2 carnitine | CD4+ T cells |
| IL17A | C18n_QI64 | tauro-alpha-muricholate/tauro-beta-muricholate | CD4+ T cells |
| LIF | HILp_QI10949 | cortisol | CD4+ T cells |
| LIF | HILp_QI11887 | C16 carnitine | B Plasma cells, CD4+ T cells |
| LIF | HILp_QI12494 | C18:1 carnitine | CD4+ T cells |
| LIF | C8p_QI212 | cholesterol | CD4+ T cells |
| LIF | C8p_QI66 | C16:0 CE | CD4+ T cells |
| LIF | HILp_QI10472 | 21-deoxycortisol | CD4+ T cells |
| LIF | C8p_QI50 | C16:0 Ceramide (d18:1) | Plasmablasts |
| LIF | C18n_QI33 | 17-methylstearate | B Plasma cells |
| LIF | C8p_QI184 | C53:3 TAG | Plasmablasts |
| LIF | HILn_QI48 | docosapentaenoate | B Plasma cells |
| PF4 | HILn_QI93 | orotate | Plasmablasts |
| PF4V1 | HILp_QI8763 | sphinganine | Plasmablasts |
| PF4V1 | C8p_QI212 | cholesterol | CD4+ T cells |
| PF4V1 | C8p_QI72 | C18:1 CE | CD4+ T cells |
| PF4V1 | C8p_QI82 | C22:6 CE | CD4+ T cells |
| PF4V1 | C8p_QI50 | C16:0 Ceramide (d18:1) | Plasmablasts |
| PF4V1 | C18n_QI33 | 17-methylstearate | B Plasma cells |
| PF4V1 | C8p_QI184 | C53:3 TAG | Plasmablasts |
| PF4V1 | HILn_QI48 | docosapentaenoate | B Plasma cells |
| TGFB3 | HILn_QI93 | orotate | Plasmablasts |

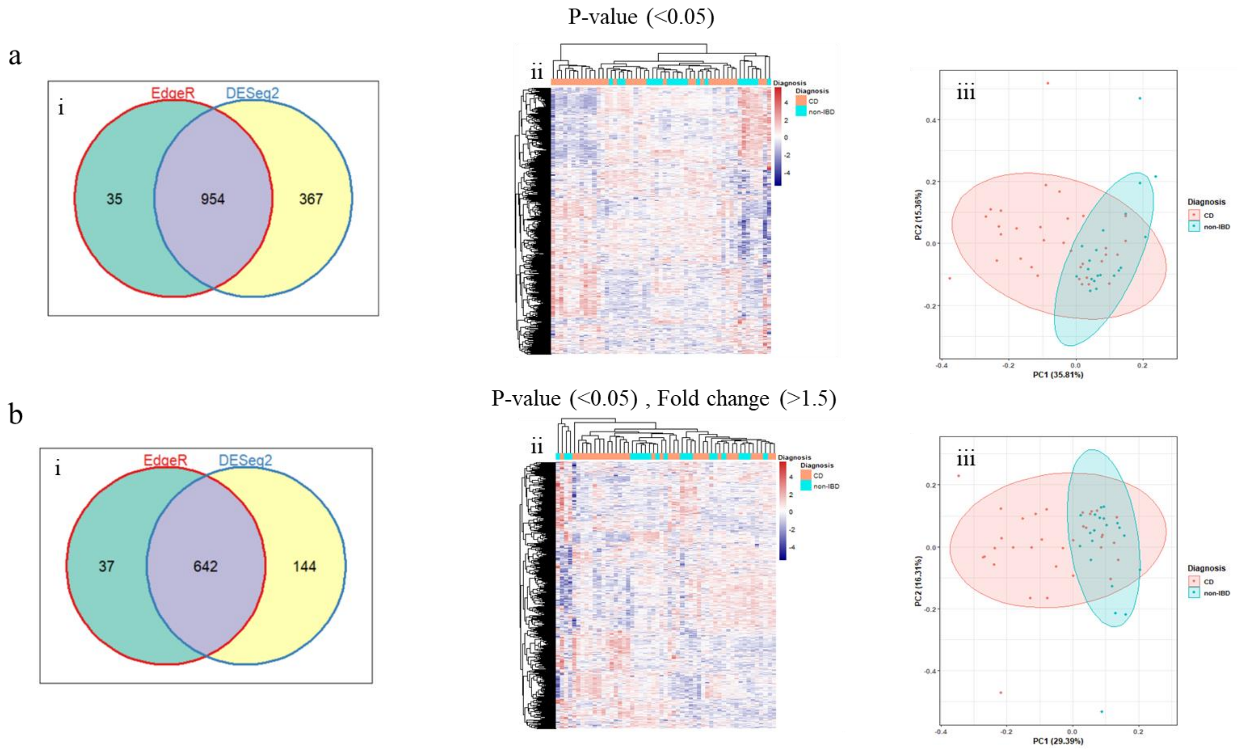

**Supplemental Figure 1, Differential expression analysis in CD in comparison with non-IBD controls. a(i-iii)**, Venn diagram, heatmap clustering, and PCA plots for significant differentially expressed genes identified by DESeq2 and edgeR in CD/control having adjusted p-value < 0.05. **b(i-iii),** Venn diagram, heatmap clustering, and PCA plots for significant differentially expressed genes identified by DESeq2 and edgeR in CD/control having adjusted p-value < 0.05 plus fold change > 1.5 and < -1/1.5.

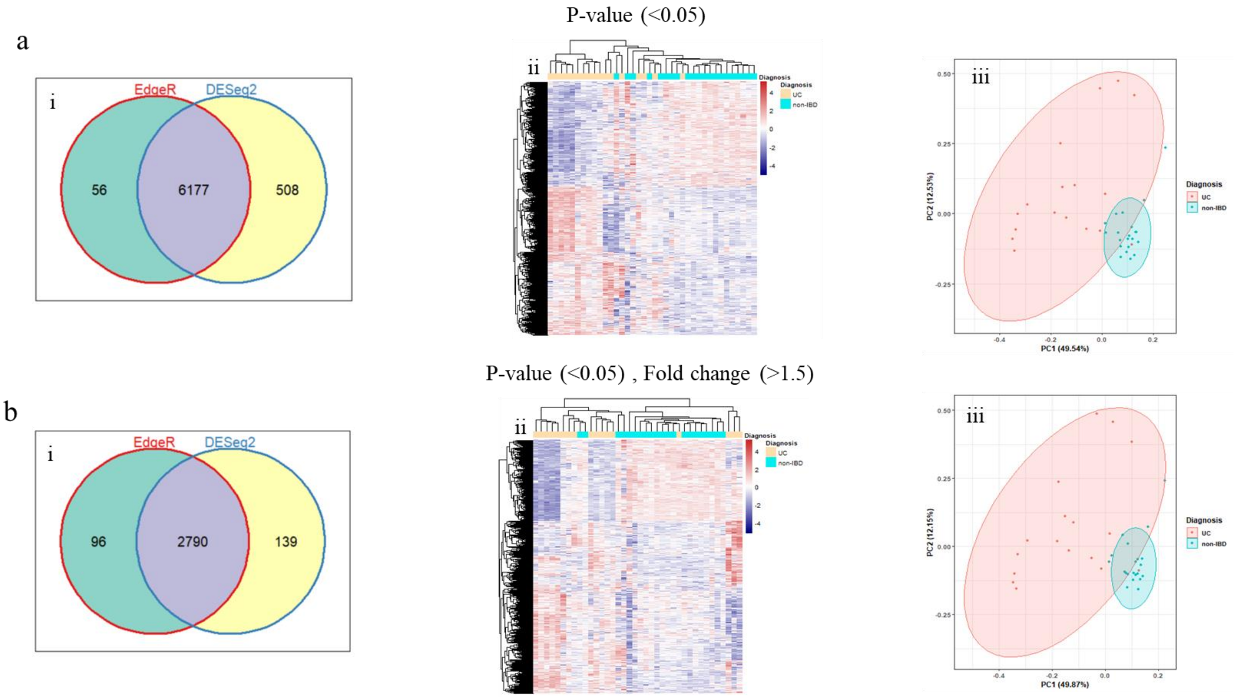

**Supplemental Figure 2, Differential expression analysis in UC in comparison with non-IBD controls**. **a(i-iii)**, Venn diagram, heatmap clustering, and PCA plot for significant differentially expressed genes identified by DESeq2 and edgeR in UC/control having adjusted p-value < 0.05. **b (i-iii)**, Venn diagram, heatmap clustering, and PCA plot for significant differentially expressed genes identified by DESeq2 and edgeR in UC/control having adjusted p-value < 0.05 plus fold change > 1.5 and < -1/1.5.

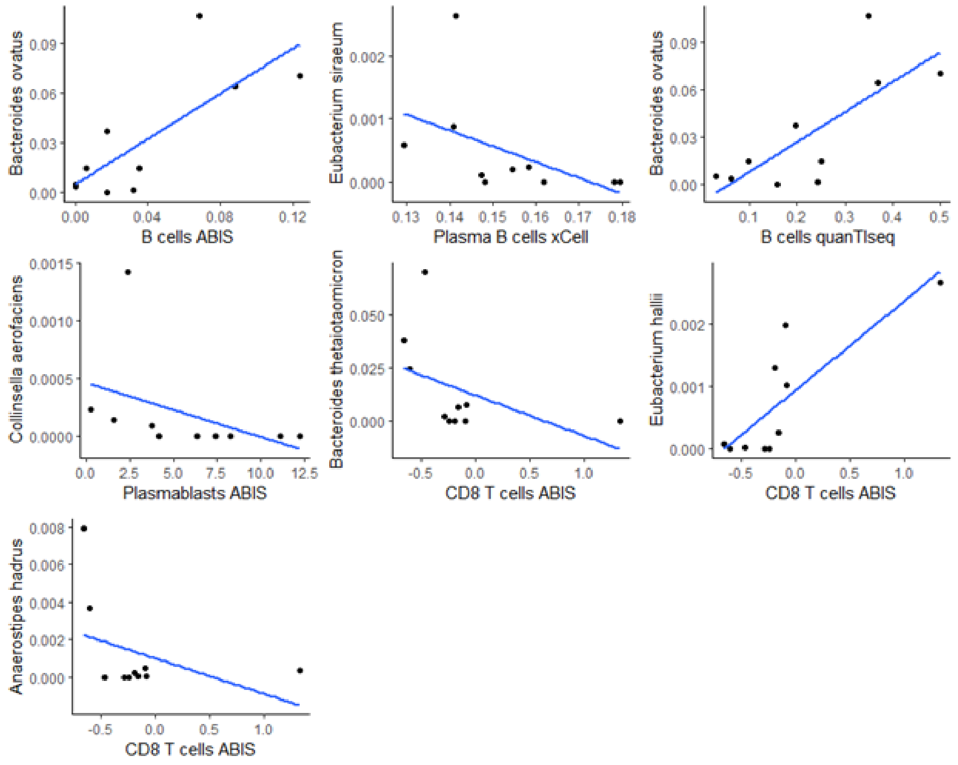

**Supplemental Figure 3, Scatter plots for significant microbe abundance versus immune cell frequency pairs for patients with active CD.**

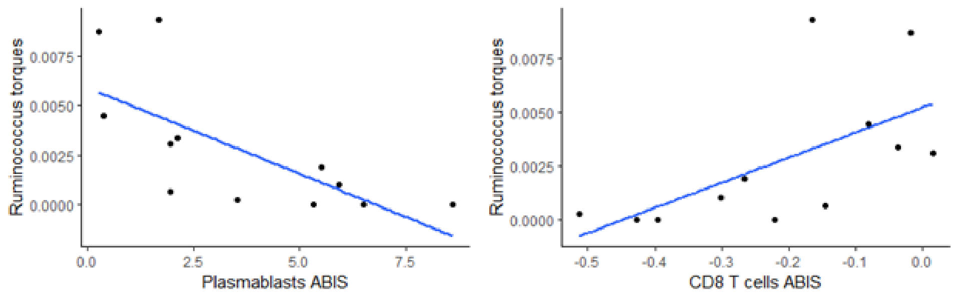

**Supplemental Figure 4, Scatter plots for significant microbe abundance versus immune cell frequency pairs for all UC patients.**

a

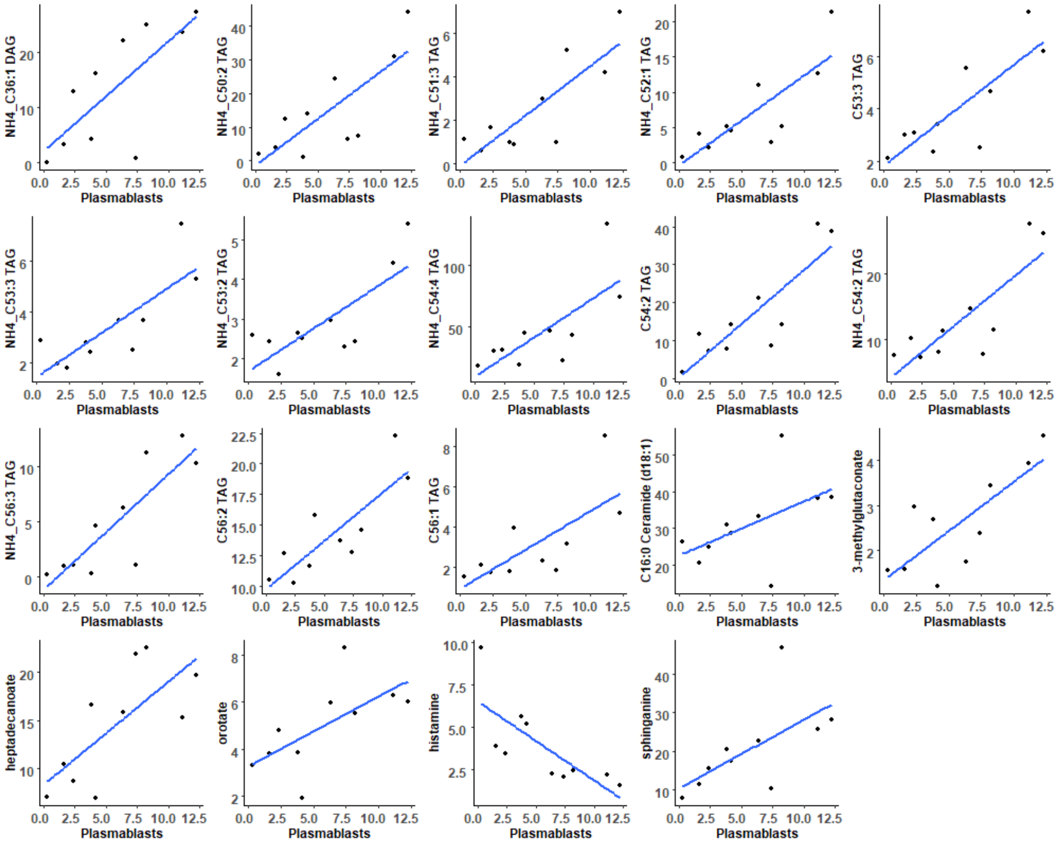
NH4_C36:1 DAG
NH4_C50:2 TAG
NH4_C51:3 TAG
NH4_C52:1 TAG
C53:3 TAG
NH4_C53:3 TAG
NH4_C53:2 TAG
NH4_C54:4 TAG
C54:2 TAG
NH4_C54:2 TAG
NH4_C56:3 TAG
C56:2 TAG
C56:1 TAG
C16:0 Ceramide (d18:1)
3-methylglutaconate
heptadecanoate
orotate
histamine
sphinganine
Plasmablasts

b

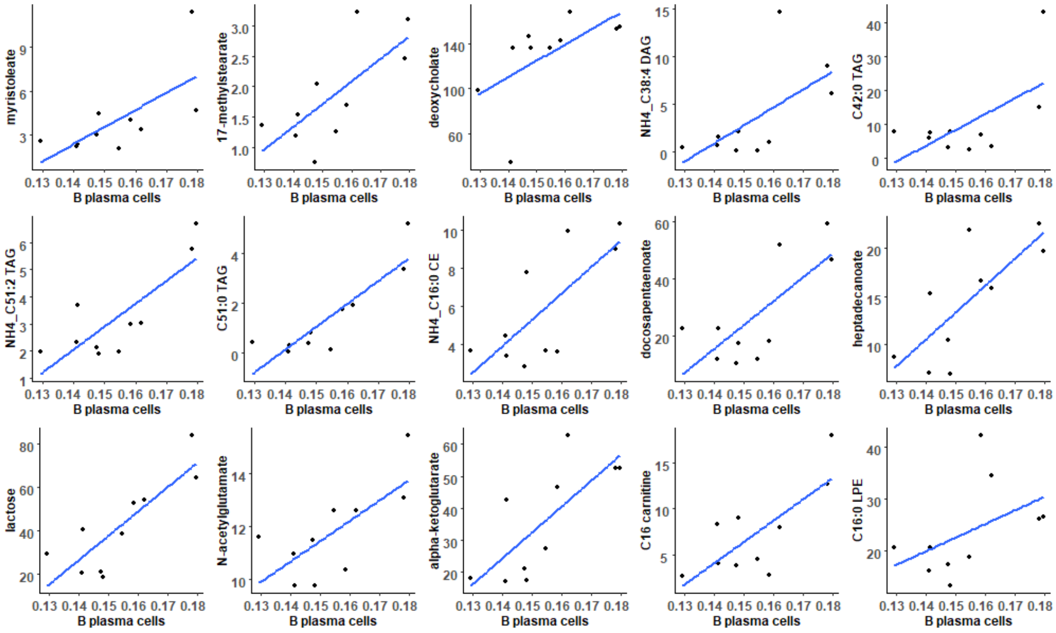
myristoleate
17-methylstearate
deoxycholate
NH4_C38:4 DAG
C42:0 TAG
NH4_C51:2 TAG
C51:0 TAG
NH4_C16:0 CE
docosapentaenoate
heptadecanoate
lactose
N-acetylglutamate
alpha-ketoglutarate
C16 carnitine
C16:0 LPE
B plasma cells

c

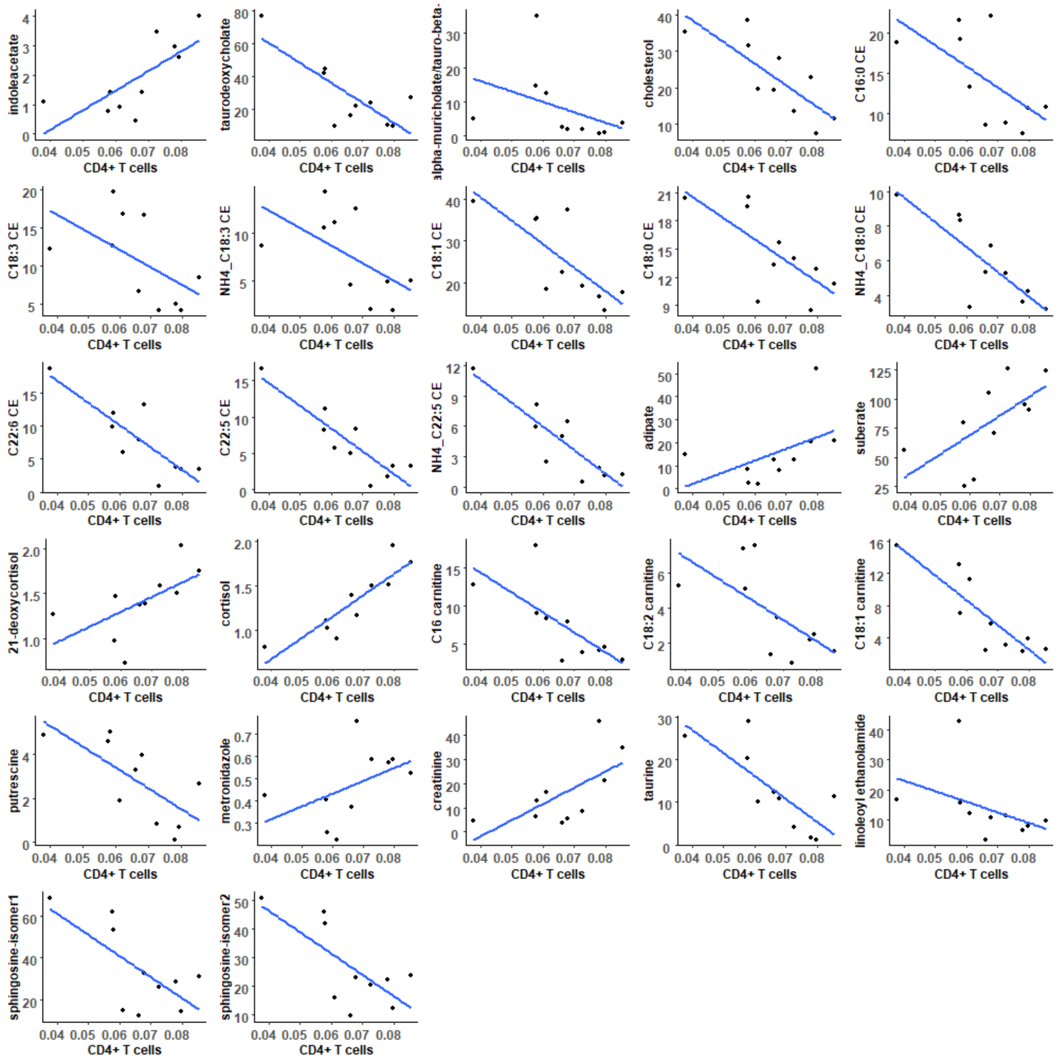

**Supplemental Figure 5, Metabolome and immune cell type association for ileum active CD (a-c).**

Scatter plots to show the association of significant metabolites with different predicted immune cell type fractions. The within-method sum normalized peak intensities of metabolites converted to sqrt(ppm) for visualization purposes.
